# Incentive valence differentially engages open- and closed-loop basal ganglia circuits during movement initiation

**DOI:** 10.64898/2025.12.21.695842

**Authors:** Neil M. Dundon, Elizabeth J. Rizor, Joanne Stasiak, Jingyi Wang, Taylor Li, Kiana Sabugo, Christina Villanueva, Parker Barandon, Viktoriya Babenko, Renee Beverly-Aylwin, Alexandra Stump, Tyler Santander, Andreea C. Bostan, Regina C. Lapate, Scott T. Grafton

**Affiliations:** Department of Psychological & Brain Sciences, University of California, Santa Barbara, Santa Barbara, CA 93106, USA; Aligning Science Across Parkinson’s (ASAP) Collaborative Research Network, Chevy Chase, MD 20815, USA; Department of Social and Psychological Sciences, University of Huddersfield, Huddersfield HD1 3DH, United Kingdom; Department of Neuroscience, University of California, Berkeley, Berkeley, CA 94720, USA; Center for Cognitive Neuroscience, Duke University, Durham, NC 27708, USA; BIOPAC Systems, Inc, Goleta, CA 93117, USA; University of Arizona College of Medicine, Tucson, AZ 85724, USA; Department of Neurobiology, University of Pittsburgh School of Medicine, Pittsburgh, PA 15213, USA

**Author notes:** Corresponding author: Neil M. Dundon, Department of Psychological & Brain Sciences, University of California, Santa Barbara, Santa Barbara, CA 93106, USA. These authors contributed equally to this work (Neil M. Dundon, Elizabeth J. Rizor). Preprint: A preprint of this manuscript will be posted on bioRxiv no later than journal submission. Classification: Major: Biological Sciences Minor: Neuroscience.

**Keywords:** motivation, motor control, striatum, functional connectivity, Parkinson’s disease

## Abstract

Incentives modulate voluntary movement, yet the circuitry channeling these signals into motor output remains unclear. Classical models emphasize a closed-loop circuit (CLC) linking dorsal putamen (PUTd) with motor cortex, but this pathway is anatomically segregated from affective processing regions. Anatomical and clinical evidence point to an alternative: an open-loop circuit (OLC) from ventral putamen (PUTv) that may route affective signals to motor cortex. Here, we conducted two experiments to test whether a functional OLC exists in humans and whether it is differentially engaged by incentive conditions. First, in 7 T resting-state fMRI (multi-echo), PUTv showed robust functional connectivity with both affective and motor regions, including the cingulate motor area (CMA), even after accounting for PUTd variance. This connectivity pattern supports the plausibility of an independent pathway linking affective basal ganglia regions to the motor cortex. Second, in 3 T task fMRI (incentivized reaching), jackpot (high-reward) and robber (high-loss avoidance) incentive conditions produced distinct behavioral and neural signatures. Jackpot produced a speed–accuracy trade-off, with faster movement initiation but more false starts. Neurally, this coincided with engagement (BOLD responses relevant for initiation speed) being reduced in CLC nodes but not in OLC. Robber, in contrast, eliminated engagement in both OLC and CLC nodes, instead recruiting stopping-related regions (e.g., STN), consistent with an avoidance phenomenology. Together, these findings support a versatile architecture for movement initiation that flexibly engages distinct cortico-subcortical circuits depending on incentive phenomenology, and offer a candidate mechanism through which affective salience and valence modulate voluntary movement.

**Significance Statement:** Affective signals profoundly influence movement, yet the mechanisms linking motivationally relevant contexts with motor behavior remain unclear. Combining ultra-high field (7 T) connectomics with task-based (3 T) neuroimaging, we provide the first systems-level evidence in humans for such a mechanism: a ventral putamen-centered open-loop circuit (OLC) connecting affective and motor areas, operating alongside the canonical dorsal putamen-centered closed-loop sensorimotor circuit (CLC). Critically, the phenomenological quality of incentive (how it is construed as reward versus threat) rather than magnitude alone, likely determines which circuit dominates during movement initiation. These findings help to explain paradoxical kinesia in Parkinson’s disease, where affective contexts can bypass degraded sensorimotor circuits, and establish foundations for context-based therapeutic interventions.

## Introduction

Control and execution of voluntary movement is sensitive to a host of contextual factors including affective and reward driven incentive (1–5). Both cortical and subcortical regions, including the basal ganglia and amygdala, play a role in processing affective and reward information, suggesting an indirect influence over motor cortical regions (6). However, the mechanisms by which affective signals are integrated into movement remain unclear.

Traditional neuroanatomical understanding of basal ganglia interactions with the motor cortex (7,8) in mammals centers on a canonical closed-loop circuit (CLC) entering the basal ganglia via the dorsal (“sensorimotor”) putamen (PUTd), projecting through globus pallidus internus (GPi), and returning to primary motor cortex (M1) via ventrolateral thalamus (VL). This sensorimotor CLC is anatomically and functionally distinct from other closed-loop circuits that link the basal ganglia with cognitive and affective regions of the cerebral cortex (7). The CLC plays a crucial role in voluntary movement, as underscored by the impairments following loss of dopaminergic innervation of PUTd (9) in Parkinson’s disease (PD), which include slowness and difficulty in initiating movement (10).

Yet, clinical observations suggest that while central, the CLC may not be the sole pathway supporting movement initiation. For instance, although the CLC is impaired in PD, affectively salient events such as car accidents (11), natural disasters (12, 13) and incentives (14) can transiently normalize movement initiation (15), suggesting alternative pathways through which affective signals might influence movement initiation in motivationally relevant contexts. One candidate circuit has been proposed by results from neuroanatomical tracing studies in nonhuman primates (NHPs). Retrograde transneuronal transport of rabies virus from injections in the primary motor cortex (M1) in NHPs shows that both PUTd and the ventral (“limbic”) putamen (PUTv) send multi-synaptic projections to M1. Notably, while PUTd is a target of M1 input, PUTv does not receive inputs from M1, but from the amygdala and cortical areas involved in affective and incentive based functions (16,17). The non-canonical connectivity profile defines a putative open-loop circuit (OLC; Fig. 1A) that may allow PUTv and related interconnected areas involved in affective functions to influence motor output (18–21). PUTv also shows spared dopaminergic innervation in primate models of PD (22). In humans with PD, PET imaging similarly shows that the ventral putamen is relatively spared, with the ventrorostral region remaining least affected even in advanced disease (23).

**Fig. 1.**
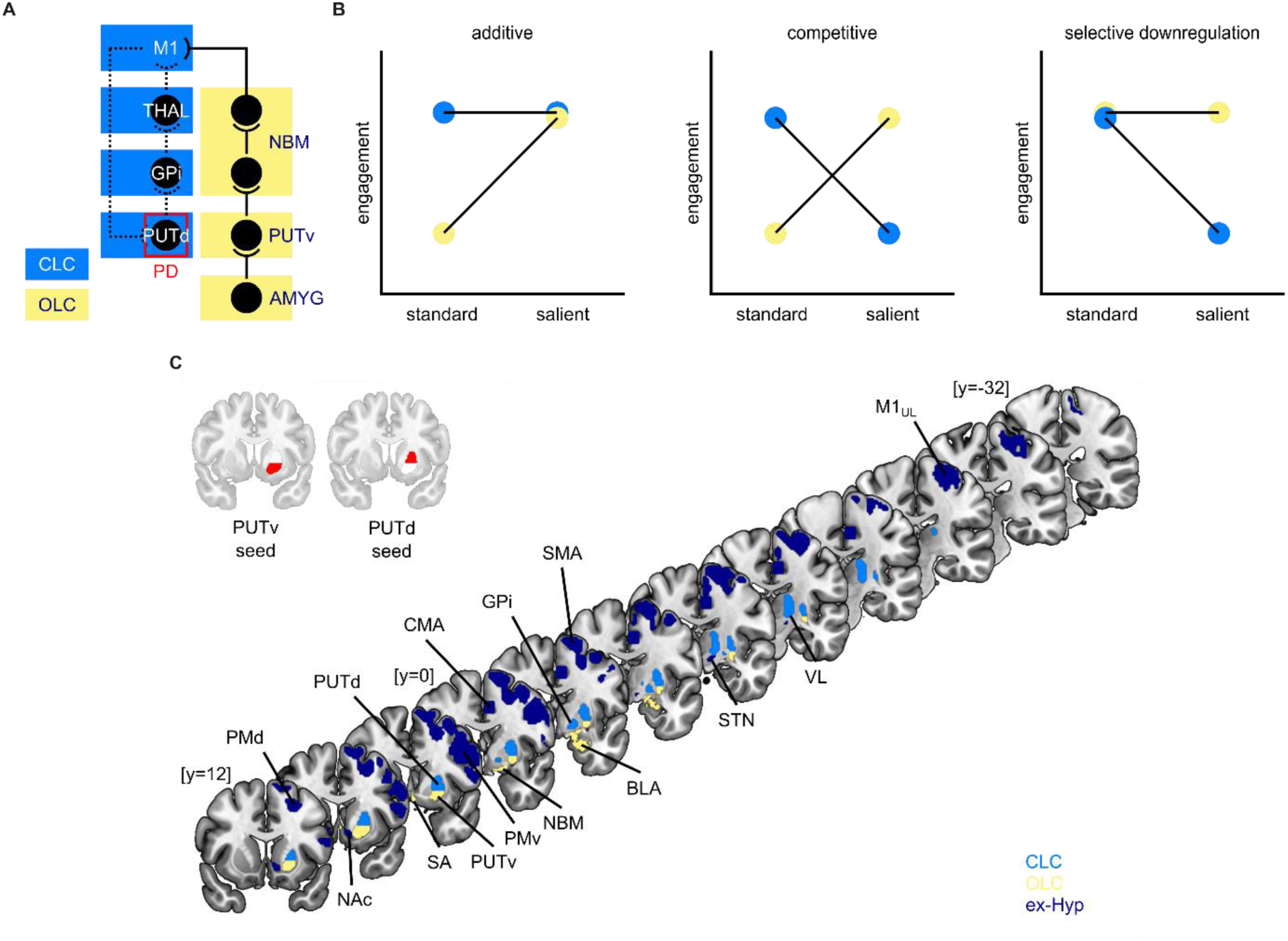
(A) Canonical closed-loop circuit (CLC) and putative open-loop circuit (OLC), with site of disruption in Parkinson’s disease (PD). (B) Three hypothesized patterns of circuit engagement by incentive salience: additive, competitive, and selective downregulation. (C) Anatomical regions of interest used across experiments, comprising CLC and OLC nodes, plus regions external to our core hypotheses, termed ex-hypothesis (ex-Hyp) nodes, that include motor and reward areas. Anatomical abbreviations are defined in Table 1 in SI.

Combined with the clinical observations in PD, these anatomical and imaging findings suggest that PUTv, and its embedding in the putative OLC, might afford affective or incentive based processes an influence over voluntary movement under motivationally relevant conditions. However, two critical questions remain unanswered: (1) does the healthy human brain exhibit a functional organization consistent with a PUTv-centered OLC that couples affective and motor systems, independently of PUTd? And (2) can activity in nodes of the putative OLC (particularly PUTv) support movement initiation under varying incentive conditions, and are the CLC and OLC differentially engaged across these conditions?

Prior systems-neuroscience experiments in humans offer key hypotheses about how incentive conditions might modulate control via differential engagement of OLC and CLC nodes. We here test three differential engagement patterns (Fig. 1B). Engagement may be *additive*, where incentives recruit OLC alongside CLC to jointly facilitate movement initiation. Such additive recruitment is consistent with cost–benefit frameworks of movement vigor (24), where value and vigor nodes converge on motor effectors to scale vigor according to expected utility. Engagement may further be *competitive*, with incentives recruiting OLC while suppressing CLC, leading to a shift in control. Such a “winner-take-all” pattern is seen during ventral (value) to dorsal (habit) striatal control shift in human addiction (25) and is similarly observed in rodents, where threat drives defensive circuits to dominate and suppress feeding, in contrast to dominant feeding circuits in the absence of threat (26). Alternatively, engagement may reflect *selective downregulation*, where both circuits are typically engaged, but high incentive salience disengages some circuitry in order to optimize for parsimony under immediate demands. Conventionally, this is observed when incentives prune cognitive control networks, increasing speed at the cost of task accuracy (27–29). Importantly, selective downregulation of hyperactive nodes can also aid neurorehabilitation (30,31), consistent with models of bradykinesia in PD that implicate excessive activity in canonical stopping circuits (32).

Thus, to identify the putative OLC in humans and examine its possible role in movement initiation, we conducted two complementary neuroimaging studies. First, we examined the plausibility of a PUTv-centred OLC using ultra-high field (7 T) multi-echo resting-state fMRI, leveraging this paradigm’s spatial-resolution and signal-to-noise advantages to assess whether PUTv is reliably functionally connected with both affective and motor regions, distinct from PUTd. Second, we analyzed task-based fMRI from a separate 3 T sample performing an incentivized reaching task under three incentive conditions (high-reward, high-loss, and standard-reward) that manipulated both salience and valence.

Using state-of-the-art BOLD signal modeling (33), we examined how activation duration in predefined CLC and OLC nodes (Fig. 1C) scaled with movement initiation speed, reasoning that circuits facilitating initiation should show the strongest responses during the fastest movements. We then tested which engagement pattern (Fig. 1B) best explained how salience (high-reward vs standard-reward) and valence (high-reward vs high-loss) shaped BOLD activation across the two circuits.

## Results

### Experiment 1

#### Connectomic evidence of PUTv connections with affective and motor regions

To test the plausibility of a putative PUTv-centered open-loop circuit (OLC), twenty-eight healthy human participants (13 female, mean age 29.68±4.78 years) completed a high-resolution resting-state fMRI scan at 7 T. To isolate intrinsic functional connectivity, we combined multi-echo fMRI with physiological monitoring and ICA denoising to selectively retain BOLD signal (see Methods).

Using PUTv and PUTd seeds, we compared their respective connectivity profiles with anatomically-defined affective and motor targets (Fig. 1). Our analysis targets included: (1) motor cortical areas, (2) established sensorimotor CLC nodes (GPi and VL), and (3) putative OLC nodes interconnected with PUTv (basal forebrain and amygdala).

We found that PUTv in the left hemisphere exhibited significant functional connectivity with multiple motor (upper limb area of motor cortex (M1_UL_), supplementary motor area (SMA), cingulate motor area (CMA)) and affective (nucleus basalis of Meynert (NBM), basolateral amygdala (BLA), central nucleus of the amygdala (CeA)) regions (summarized in the connectome plot in Fig. 2A; all p <0.05, FDR-corrected). All connections were bilateral except for BLA and M1_UL_, which were ipsilateral. PUTv was also functionally connected bilaterally with the nucleus accumbens (NAc) and ipsilaterally with VL. A similar connectivity profile was observed with a PUTv seed in the right hemisphere (Fig. S2). In contrast, PUTd was preferentially connected with motor regions (CMA, SMA, dorsal premotor cortex (PMd), M1_UL_, ventral premotor cortex (PMv)), showing connectivity with fewer affective regions (NBM, CeA) (Fig. 2B; all p <0.05, FDR-corrected). NBM connections were bilateral, CeA was unilateral. Meanwhile, CMA, PMd and M1_UL_ were ipsilateral and remaining motor targets bilateral. PUTd was also functionally connected ipsilaterally with VL (left). A similar connectivity profile was observed with a PUTd seed in the right hemisphere (Fig. S3).

**Fig. 2.**
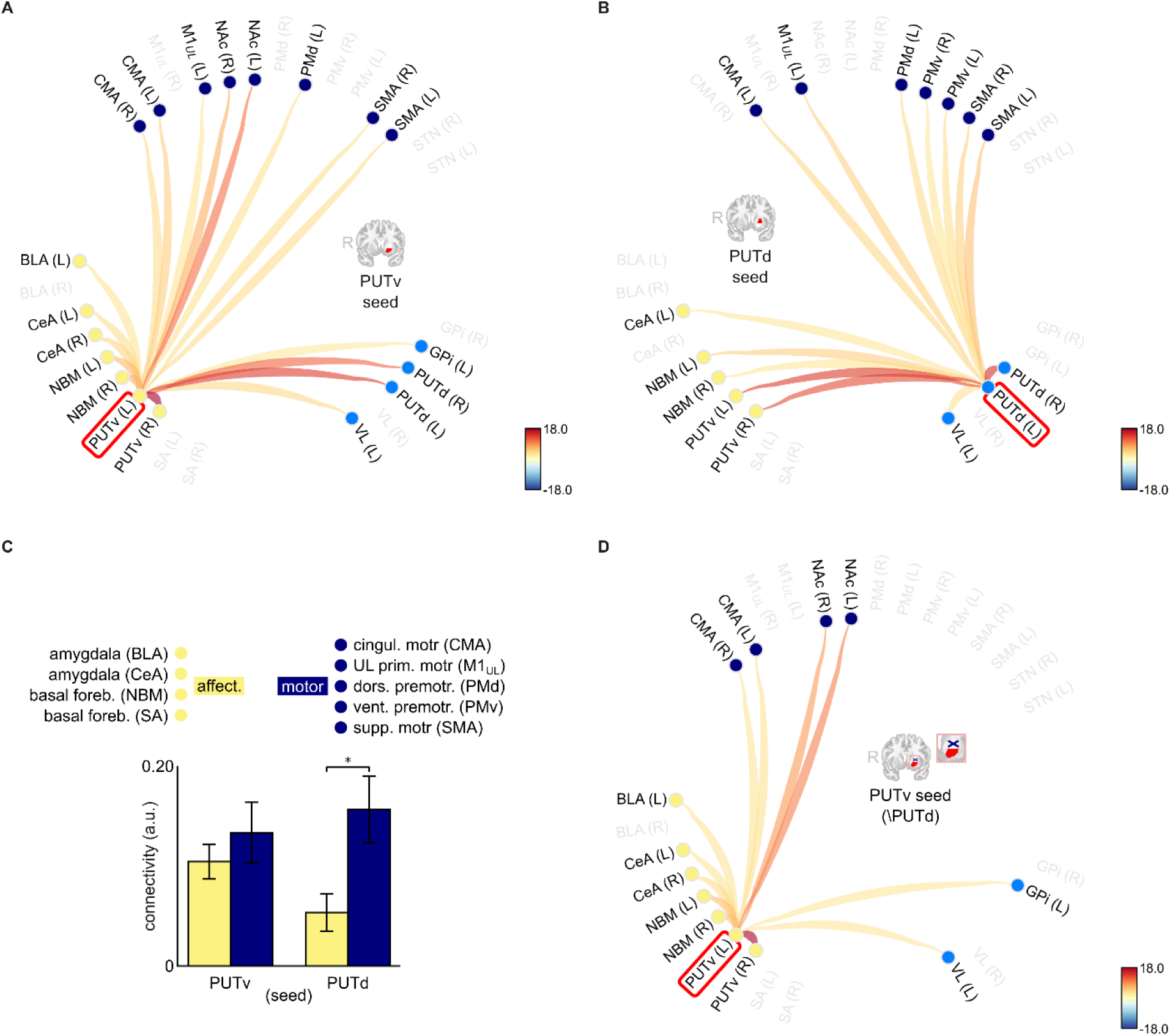
(A) Left PUTv connectivity with OLC (light blue), CLC (yellow), motor and other regions (dark blue). (B) Left PUTd connectivity. (C) Interaction between putamen subregion seed and target network. (D) Left PUTv connectivity after controlling for shared variance with left PUTd. Connection strengths in A, B, and D represent correlation t-statistics.

To formally evaluate whether dorsal and ventral putamen subregions differed systematically in their functional connectivity with motor versus affective networks, we tested for region-by-network differences using an ANOVA (Fig. 2C). This revealed a significant interaction between putamen subregion and target network (F(1,27) = 8.45, *p* = 0.007, η² = 0.019). Post-hoc comparisons indicated that PUTd was selectively coupled to motor targets (p = 0.03, Bonferroni corrected, *d* = –0.72), whereas PUTv was functionally connected with both motor and affective regions to a similar degree, a profile potentially capable of linking the two. Thus, PUTd showed motor-selective connectivity, whereas PUTv showed both motor and affective connectivity.

Finally, to test whether PUTv connectivity (in particular, its coupling with motor regions) simply reflected shared variance with PUTd, we repeated the connectivity analysis after partialling out the ipsilateral PUTd time series. We replicated our results after imposing this additional constraint, whereby left PUTv remained functionally connected to key affective regions (BLA, CeA, NBM) as well as the motor-associated CMA (all *p*< 0.05; FDR-corrected; Fig 2D), indicating integration with affective and, critically, motor circuits independently of PUTd. (Similar results were observed with this partial correlation using method, using a PUTv seed in the right hemisphere; Fig. S4). Two additional control analyses further clarified the specificity of these connectivity patterns. First, partialling out PUTv variance eliminated the PUTd-CMA association (SI Figure S5, S6), suggesting that apparent PUTd–CMA connectivity is itself dependent on shared variance with PUTv. Second, the PUTv–CMA association remained robust when controlling for amygdala variance (SI Figure S7-S10), indicating that PUTv–CMA connectivity is independent of common amygdala inputs.

Collectively, these findings demonstrate that PUTv exhibits independent functional connectivity with both motor and affective networks, consistent with the proposed OLC architecture whereby PUTv provides a pathway through which affective and incentive signals influence motor output in healthy humans.

### Experiment 2

#### Task fMRI evidence that incentive conditions modulate movement initiation and circuit engagement

To examine how varying incentive conditions modulate movement initiation and movement-related BOLD activity in CLC versus OLC nodes, sixty-eight healthy human participants (50 female, mean age 20.75±1.86 years) performed a speeded precision joystick reaching task during an fMRI scan at 3T. Each trial offered a jackpot ($1.60 high-reward), robber ($1.60 high-loss avoidance) or standard ($0.20 reward) incentive (Fig. 3A). Success was defined as the cursor reaching and being-held at the cued target before a preset group deadline, which determined if incentives were met. The task comprised 300 trials in total (10% jackpot, 10% robber, 80% standard), with participants achieving 56.8% success overall.

**Fig. 3.**
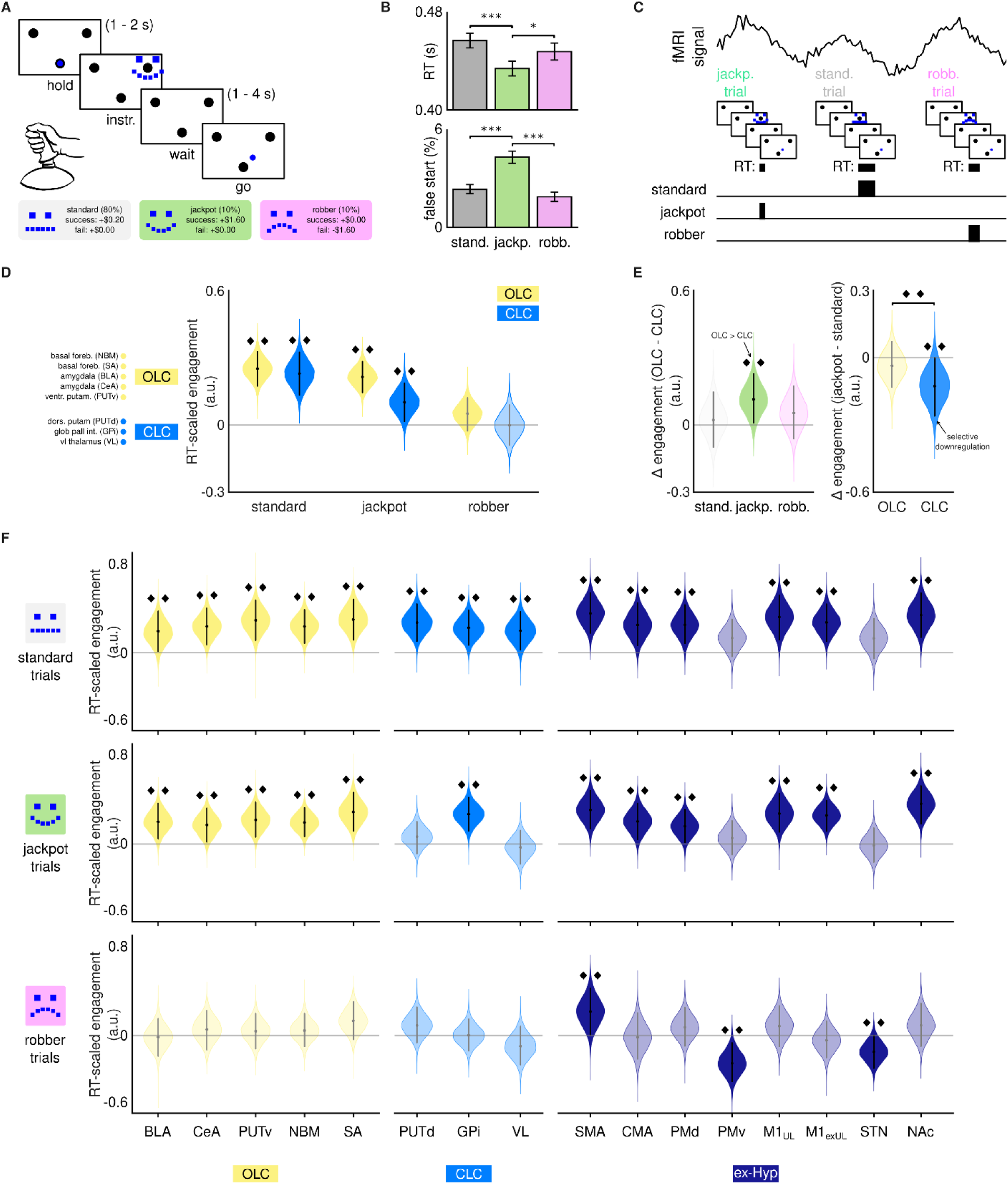
(A) Task design with jackpot, robber, and standard reward conditions. (B) Median movement initiation times (RT) and average false starts (movement prior to “go” cue) by incentive condition; error bars show SEM. (C) RT-scaled BOLD modeling approach. (D) Circuit-level engagement across conditions: both OLC and CLC engaged under standard reward, but neither under robber. (E) Relative circuit engagement across conditions and relative jackpot engagement across circuits, combine to demonstrate selective downregulation of CLC. (F) Node-level analysis for OLC, CLC and ex-hypothesis nodes across conditions. Violin plots (D-F) show Bayesian posterior distributions of the location parameter μ, where positive/negative values indicate greater BOLD response with shorter/longer RT. Dots indicate posterior mean, lines span HDI. *p<.05; ***p<.001; ♦♦ indicates a Bayesian credible effect: the posterior HDI for engagement excludes 0, or the HDI of the difference excludes 0 for pairwise comparisons.

We first assessed behavioral effects of incentive conditions (all kinematic variables in SI Figure 1, SI Table 3). While success rates showed an overall effect of incentive (F(2,134)=4.16, p=0.018), pairwise comparisons were non-significant (all p_fdr_ >0.087). Critically, incentives robustly modulated movement initiation speed (RT; Fig. 3B; F(2,134)=18.5, p<0.001), with participants initiating reaches faster in the jackpot condition (440ms) compared to both standard (460ms; p_fdr_ <0.001) and robber (450ms; p_fdr_ = 0.013). Beyond initiation, the robber condition uniquely increased peak velocity (SI Figure 1, SI Table 3). Strikingly, the jackpot condition uniquely increased false starts, that is, movements initiated prior to the go-cue (Fig. 3B; F(2,134) = 25.4; p<0.001). This pattern suggests the jackpot condition prioritized initiation speed at the cost of accuracy, whereas robber modulated a different kinematic phase (execution velocity).

#### RT-scaled BOLD activation reveals selective downregulation of CLC under positive incentive

We next tested whether CLC and putative OLC nodes showed differential BOLD responses associated with movement initiation across incentive conditions. For this, we examined how BOLD signal duration in predefined CLC and OLC nodes (Fig. 1C) scaled with movement initiation speed. To contextualize these circuits’ effects within the broader motor and reward system, we additionally included regions external to our core hypotheses, termed ex-hypothesis (in Fig. 1, 3) regions (SMA, CMA, PMd, PMv, M1_UL_, subthalamic nucleus (STN), and NAc) in the model. We employed an extension of recent BOLD signal modeling approaches (33) using condition-specific RT-scaled regressors at movement initiation (see Methods, Fig. 3C). Using a hierarchical Bayesian approach, we estimated RT-scaled BOLD activation at the group level for each node and condition (𝜇_𝑐𝑜𝑛𝑑𝑖𝑡𝑖𝑜𝑛,𝑟𝑒𝑔𝑖𝑜𝑛_), which were then averaged across nodes at each posterior draw to derive circuit-level RT-scaled estimates (𝜇_𝑐𝑜𝑛𝑑𝑖𝑡𝑖𝑜𝑛,𝑐𝑖𝑟𝑐𝑢𝑖𝑡_) (see Methods, Fig. 3D). We reasoned that circuits facilitating movement initiation should show greater BOLD responses during faster movements. Accordingly, positive μ values indicate what we operationally define as *engagement*, i.e., greater BOLD response with faster movement initiation.

First, we examined OLC and CLC’s *overall* engagement in the three incentive conditions (Fig. 3D). In the standard condition, nodes of both circuits showed credible engagement, with posterior HDIs excluding zero in each case (𝔼(𝜇*_standard,CLC_*)= 0.251, HDI = [0.175,0.330]; 𝔼(𝜇*_standard,CLC_*) = 0.229, HDI = [0.130, 0.322]). This pattern replicated under jackpot (𝔼(𝜇*_jackpot,OLC_*) = 0.215, HDI = [0.146, 0.284]; 𝔼(𝜇*_jackpot,OLC_*)= 0.101, HDI = [0.013, 0.184]), albeit with weaker engagement in CLC (see below). In contrast, in the robber condition, neither circuit showed credible engagement, with HDIs including zero (𝔼(𝜇*_robber,OLC_*) = 0.050, HDI = [-0.025 , 0.122]; 𝔼(𝜇*_robber,OLC_*) = - 0.002, HDI = [-0.094, 0.089]). Thus, nodes of both CLC and OLC are robustly engaged (greater BOLD response with faster RT) when movements are motivated by positive incentive (standard or jackpot). However, this engagement is notably absent in the robber condition, consistent with robber not eliciting faster movement initiation behaviorally. This dissociation suggests valence influences OLC and CLC involvement in movement initiation.

Next, we examined OLC and CLC’s *relative* engagement in the three incentive conditions, to probe for shifts in the balance of control (Fig. 3E). For this, we directly contrasted engagement across circuits (Δ*_condition_*= 𝜇*_condition, OLC_* − 𝜇*_condition, CLC_*) for each condition. In the standard condition, circuits showed no credible difference in engagement (𝔼(Δ*_standard_*) = 0.021, 89% HDI = [-0.102, 0.144]), suggesting balanced engagement during movement motivated by standard reward. Similarly, under robber, no credible difference emerged between circuits (𝔼(Δ*_robber_*) = 0.052, HDI = [-0.059, 0.175]), though taken with results above, this likely reflects mutual disengagement rather than balanced activation.

Strikingly, in the jackpot condition, OLC nodes showed credibly stronger engagement than CLC (𝔼(Δ_𝑗𝑎𝑐𝑘𝑝𝑜𝑡_) = 0.113, HDI = [0.004, 0.224]). To directly test whether this constitutes selective downregulation of CLC, independent of dynamics in OLC, we examined within-circuit changes from standard to jackpot conditions (Δ*_circuit_*= *𝜇_jackpot,circuit_* − 𝜇*_standard,circuit_*) for each circuit (Fig. 3E). This revealed that while OLC nodes maintained consistent engagement across both conditions (𝔼(Δ*_OLC_*) = -0.036, HDI = [-0.141, 0.068]), CLC showed credible downregulation under jackpot 𝔼(Δ*_CLC_*) = - 0.128, HDI = [-0.256, -0.002]).

To pinpoint the sources of the apparent CLC downregulation, we next examined engagement at the level of individual nodes (Fig. 3F). Consistent with stable engagement, all OLC nodes were credibly engaged in both standard and jackpot conditions (0 ∉ all posterior HDIs; Fig. 3F, left panels, upper and middle row). In contrast, within the CLC (middle panels), a selective shift emerged: all nodes were engaged in the standard condition (0 ∉ all posterior HDIs; Fig. 3F, middle panels, upper row), but under jackpot, only GPi remained engaged (𝔼(𝜇_𝐺𝑃𝑖,𝑗𝑎𝑐𝑘𝑝𝑜𝑡_) = 0.268, HDI = [0.118, 0.418]; Fig. 3F). Both VL and PUTd no longer showed credible engagement (0 ∈ each posterior HDI; (𝔼(𝜇_𝑉𝐿,𝑗𝑎𝑐𝑘𝑝𝑜𝑡_) = -0.030, HDI = [-0.180, 0.122]); 𝔼(𝜇_𝑃𝑈𝑇𝑑,𝑗𝑎𝑐𝑘𝑝𝑜𝑡_) = 0.066, HDI = [-0.087, 0.201]; middle panels, middle row), implicating their downregulation as primary contributors to reduced CLC engagement in the jackpot condition.

The node-level results (Fig. 3F) further reveal that the jackpot-driven engagement shifts (relative to standard) were subcortically selective. Beyond our a priori CLC and OLC nodes, we included non-hypothesized regions in our hierarchical Bayesian model (“ex-Hyp” in Figs. 1, 3) to contextualize circuit-level effects within the broader motor system. Engagement patterns for these exploratory regions are shown in the right panels of Fig. 3F. A striking dissociation emerged: whereas CLC nodes (PUTd, VL) showed reduced engagement from standard to jackpot, all motor-cortical regions (SMA, CMA, PMd, PMv, M1_UL_, Motor cortex excluding upper limb region (M1_exUL_)) maintained stable engagement in both conditions. With the exception of PMv, all cortical regions showed credible engagement under both standard and jackpot. Among subcortical regions outside our hypothesized circuits, NAc showed credible engagement in both standard and jackpot conditions, whereas STN did not show credible engagement in either. This pattern indicates that jackpot-driven downregulation specifically targets nodes in the CLC rather than reflecting a general motor or reward system reconfiguration.

Even more strikingly, the robber condition elicited a dramatic cortical reconfiguration. Unlike the stability observed across positive-incentive conditions (i.e., standard and jackpot), most motor-cortical nodes did not show credible engagement under robber (Fig. 3F, right panel, bottom row), mirroring the subcortical disengagement observed in this condition (i.e., nodes of both OLC and CLC; Fig. 3F, left and middle panels, bottom row). SMA emerged as the sole region (cortical or subcortical) maintaining credible engagement across all three incentive conditions, suggesting a condition-invariant role in movement initiation. Remarkably, PMv and STN (neither of which were credibly engaged under standard or jackpot) uniquely exhibited negative engagement under robber. Negative engagement reflects greater BOLD response for slower RTs, i.e., the opposite pattern from nodes engaged in the two reward conditions. This inverse, valence-specific recruitment of PMv and STN may therefore implicate the robber condition (i.e., loss-avoidance) in uniquely engaging stopping (i.e., inhibitory or hesitation) processes that remain dormant during the reward-motivated movement in the other two conditions.

Collectively, these findings reveal incentive-specific engagement of CLC, OLC and cortical motor areas during movement initiation. In the standard condition, both OLC and CLC showed robust engagement (greater BOLD response with faster RT), reflecting coordinated subcortical contributions to movement initiation. High-reward (jackpot) incentive selectively downregulated CLC (particularly PUTd and VL) while preserving OLC engagement, with this subcortical reconfiguration occurring against stable motor-cortical engagement. By contrast, loss-avoidance (robber) incentive eliminated credible engagement in both subcortical circuits, reduced engagement across most motor-cortical regions (excepting SMA), and produced negative engagement (greater BOLD response with slower RT) in regions associated with stopping (PMv and STN). These divergent patterns point to a versatile architecture for movement initiation that flexibly engages distinct cortico-subcortical circuits depending on both the salience and valence of incentive conditions.

## Discussion

Affective signals influence motor control, potentially by way of the brain maintaining multiple parallel cortico-subcortical circuits differentially engaged by motivational context. Neuroanatomical evidence in nonhuman primates and clinical observations in humans suggest a PUTv-centered circuit with connectivity to affective systems (OLC) may operate alongside the canonical sensorimotor circuit through dorsal putamen (CLC), but whether these circuits exist as functionally independent in humans and, critically, whether they are differentially recruited under varying incentive conditions remains unknown. We therefore conducted two complementary neuroimaging experiments.

We first report connectomic data acquired using ultra-high field multi-echo fMRI, enabling precise regional delineation and mitigation against signal dropout, particularly in the basal ganglia (34). These data reveal that PUTv maintains balanced connectivity with both affective and motor networks, while PUTd couples preferentially to motor targets. Critically, dual connectivity persisted in PUTv even after controlling for shared variance with PUTd, suggesting functional independence from the traditional motor circuit.

Within this partialled map, PUTv showed prominent connections with amygdala nuclei (central (CeA) and basolateral (BLA)), consistent with NHP tracer evidence identifying amygdala structures as inputs to PUTv (7, 8, 35). These findings complement existing observations of a dorsal/ventral striatum gradient, whereby ventral striatum is characteristically closely associated with subcortical areas relevant for affective and motivational processes (17, 36–38). We additionally observed robust associations between PUTv and basal forebrain (NBM). While this structure may offer cholinergic input to PUTv, NHP evidence also supports its role as an output target. NHP ventral striatum (combining PUTv and NAc) bypasses the canonical GPi route to motor cortex (35, 39), targeting instead ventral pallidum and NBM, whose cholinergic outputs widely target the cerebral cortex (40–42).

PUTv signal also covaried with a broad suite of motor and premotor areas, however most of these connections were eliminated when PUTd variance was removed, suggesting they likely reflected striato-motor connectivity mediated by PUTd.

Nevertheless, PUTv robustly covaried with CMA, situated on the inferior medial wall of the caudal frontal lobe. This association persisted when controlling for both PUTd and amygdala variance, positioning CMA as a candidate node through which PUTv may influence motor circuits. This finding is noteworthy because CMA’s functional properties align with post-selection affective modulation of motor function. CMA is classically considered a specialized hub for translating affective drive into action (43), and preferentially activates for self-initiated movements (44). Recent network-neuroscience evidence further suggests that it sits downstream of decision-related regions where it initiates and implements selected actions (45). The inputs to CMA further support this post-selection modulatory role: both PUTv and the basolateral amygdala encode conjunctions of incentive valence and action tendency (46–48), but do not represent detailed movement parameters. Critically, CMA’s projections to premotor spinal laminae favor global action initiation rather than fine motor control (49–51), distinguishing it from lateral motor areas involved in precise movement execution.

While CLC and OLC circuit configurations involving CMA and basal ganglia remain to be characterized in primates, these converging functional and anatomical properties position PUTv’s preserved affective connectivity and association with CMA as an alternative route through which affective signals may influence movement initiation beyond canonical sensorimotor circuits. Direct demonstration of this functional pathway awaits further investigation.

We next report behavioral and task-derived BOLD data to examine how incentive conditions modulate initiation speed and the engagement profile of OLC nodes relative to those in CLC. First, at the behavioral level, we revealed a clear asymmetry in incentive effects: high reward (jackpot) selectively expedited movement initiation relative to both standard and high-loss avoidance (robber) conditions. Concurrently, participants also showed increased false starts under jackpot, consistent with participants optimizing for speed at the cost of accuracy under high positive incentive.

Task-derived BOLD activation was then assessed to determine which engagement pattern best explained circuit dynamics across incentive conditions: additive, competitive, or selective downregulation. Modeling RT-scaled BOLD activations allowed us to infer whether circuit nodes operationally scaled their activity with initiation speed. Under standard reward conditions, OLC, CLC, and cortical-motor nodes showed comparable RT-scaled activation, suggesting balanced engagement across the brain during typical goal-directed movement. However, incentive salience and valence drove divergent patterns of engagement, with robber and jackpot conditions seemingly engaging qualitatively distinct control architectures.

The high-reward (jackpot) incentive produced a striking dissociation consistent with our selective downregulation hypothesis: OLC nodes maintained engagement (relative to the standard condition) while CLC nodes disengaged. Node-level analysis revealed this downregulation was subcortically selective, driven by reduced engagement in PUTd and VL thalamus, while motor cortical regions maintained stable engagement (CMA, SMA, PMd, M1). Behaviorally, jackpot drove faster initiation alongside more false starts, a clear speed-accuracy trade-off. This pattern suggests that high positive incentive downweights deliberative sensorimotor basal ganglia circuits, resulting in leaner corticostriatal circuitry predominantly involving PUTv and affective regions, while prioritizing speed over accuracy (27–29). Critically, CMA maintained engagement under jackpot. As discussed above, CMA’s properties favor global movement initiation over fine control. In addition, Experiment 1 established robust functional connectivity between CMA and PUTv. Thus, one parsimonious interpretation is that with reduced CLC influence, affective signaling through CMA (potentially via PUTv) may gain relative modulatory prominence in shaping rapid motor output.

By contrast, the high-loss avoidance (robber) incentive revealed a pattern consistent with our competitive shift hypothesis. In the robber condition, neither OLC nor CLC showed credible RT-scaled activation, indicating disengagement of these circuits. Instead, robber uniquely recruited PMv and STN, regions that remained dormant under both standard and jackpot. Moreover, these regions exhibited negative engagement: greater activation with slower RT, suggesting recruitment of hesitation or stopping processes that compete with movement initiation. This pattern, i.e., distinct regions with different behavioral correlates, is consistent with broader evidence from human learning and decision-making, where rewards and costs recruit qualitatively distinct mechanisms rather than encoding valence through opposing responses in shared circuits (52,53). The negative engagement in PMv is also consistent with recent evidence in PD patients, where increased cautiousness with movement appears to shift control away from striatum to cortical areas (54). Thus, high-loss avoidance appears to engage stopping processes that compete with and may even supplant the circuits supporting movement initiation under positive incentive conditions.

The differential engagement patterns across incentive conditions are consistent with a multi-level control architecture, proposed in Fig. 4. Traditional action-selection models posit that competing action plans (including stopping or hesitation) vie for motor output (55). Our findings extend this framework, suggesting that phenomenology evoked by incentive conditions determines which of several coexisting control architectures dominates. Despite jackpot and robber conditions being economically equivalent, they elicited entirely distinct behavior and neural configurations. Thus, even for a behavior as fundamental as movement initiation, the controlling architecture itself may reconfigure based on phenomenology, specifically, the asymmetric weighting of incurred losses versus forgone gains. This would constitute a flexible control that accommodates when the greatest cost is hesitation, and also when the greatest cost is haste.

**Fig. 4.**
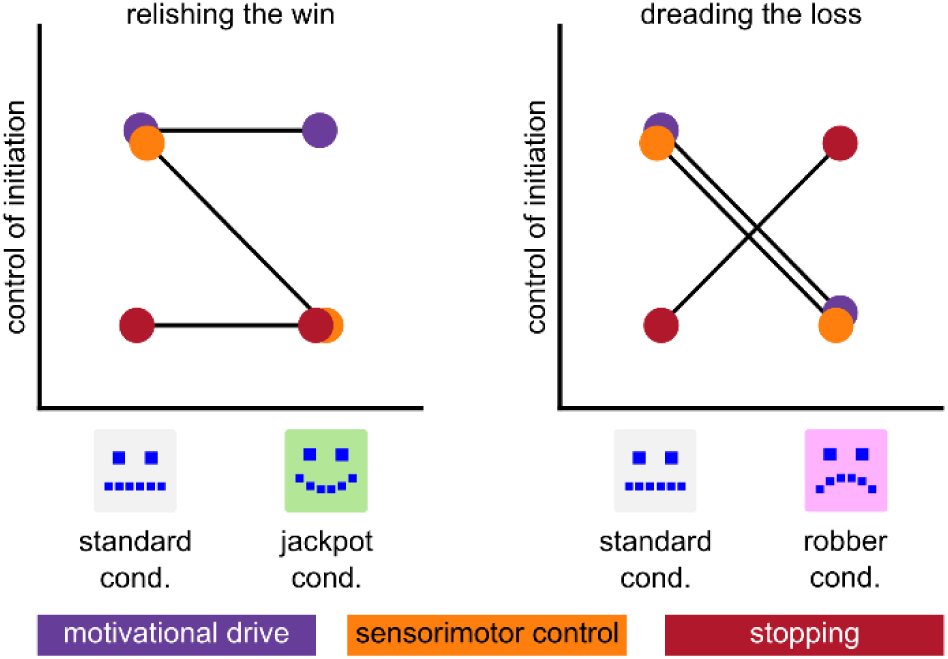
Proposed multi-level model whereby phenomenology (i.e., the subjective construal of incentive as opportunity versus threat) determines which neural architecture dominates movement initiation

Our present findings necessarily remain correlational. In the connectivity findings in particular, we cannot definitively establish that the affective regions PUTv and CMA form a directional circuit. Indeed, the co-activation patterns we observe could reflect common upstream drivers, feedback loops, or other non-causal relationships. Nevertheless, within the constraints of human neuroimaging, we advance the evidence base in specific ways. First, our ultra-high field multi-echo acquisition, signal decomposition, and partialling out of PUTd variance provide unprecedented resolution for establishing a functional integration for PUTv that is independent of nodes in canonical sensorimotor circuits. Second, we employ RT-scaled BOLD modeling that identifies which circuit nodes operationally scale their activity with initiation speed under different incentive conditions.

An important step toward greater causal inference is to translate our findings into clinical experiments, particularly in PD where convergent evidence suggests therapeutic potential. As we noted, PUTv is relatively spared in both NHP PD models (22,23) and in humans with PD, at least early in the disease. Amygdala–PUTv connectivity likewise remains preserved, at least until mid-stage disease progression (56). Moreover, the disease shows divergent trajectories across motor circuits: decreased SMA-putamen connectivity yet preserved CMA function (57, 58). Together, these findings suggest that key nodes of the proposed OLC pathway remain anatomically viable even as the canonical CLC degrades. Our model suggests that paradoxical kinesia (PK) in some Parkinson’s disease patients may occur when affectively salient contexts engage this preserved pathway, shifting control from impaired deliberative circuits. However, striatal-bypass accounts, via brainstem or cerebellum, additionally offer plausible alternative models for PK (59). A greater understanding of how the brain flexibly supports voluntary movement across affectively salient contexts may open new therapeutic avenues, from lifestyle interventions that leverage such contexts to refined deep-brain stimulation protocols.

## Materials and Methods

### Experiment 1

#### Participants and MRI Acquisition

Twenty-eight healthy, right-handed participants (15M/13F, ages 21–39) were scanned at 7T (Siemens Terra) with a 32-channel receive head coil. The protocol included T1-weighted MPRAGE anatomical imaging (0.75 mm³), multi-echo resting-state fMRI (2×2×1.5 mm, 10:52 min), and spin-echo field maps for distortion correction. Physiological signals (respiration, pulse) were recorded during scanning. All participants provided written informed consent under protocols approved by the Institutional Review Boards at University of California, Santa Barbara and the University of Southern California (USC). Full acquisition parameters and participant screening criteria are provided in SI Methods S1.

#### fMRI Preprocessing

Multi-echo fMRI data were preprocessed using ANTs, FSL, and Tedana (v24.0.1). Preprocessing included motion correction, slice-time correction, fieldmap-based distortion correction (available for 16/28 participants), and coregistration to T1 anatomy. Tedana was used to optimally combine echoes and remove non-BOLD signal components via PCA/ICA. Denoised data were normalized to MNI-152 space for connectivity analysis. Full preprocessing details are provided in SI Methods S2.

#### Functional Connectivity Analysis

Functional connectivity was computed using the CONN toolbox (v22.a). Additional confound regression included CompCor-derived anatomical noise components and bandpass filtering (0.008–0.09 Hz). Seed-to-voxel connectivity was estimated via bivariate correlation, with significance assessed using FDR correction (α = 0.05). To isolate PUTv-specific connectivity, partial correlations were computed after regressing the ipsilateral PUTd time series. Network-level analyses compared connectivity to motor (CMA, SMA, PMd, M1UL, PMv) and affective (BLA, CeA, NBM) ROIs using repeated-measures ANOVA. Full details are provided in SI Methods S3.

### Experiment 2

#### Participants and MRI Acquisition

Sixty-eight healthy participants (50F/18M, mean age = 20.75 years; SD = 1.86) were scanned at 3T (Siemens Prisma) with a 64-channel head/neck coil. The protocol included T1-weighted MPRAGE anatomical imaging (0.94 mm³), gradient-echo field maps for distortion correction, and task fMRI (2.5 mm³, TR = 1900 ms) acquired during three runs. All participants provided written informed consent under a protocol approved by the Institutional Review Board at University of California, Santa Barbara (UCSB). Full acquisition parameters and participant screening criteria are provided in SI Methods S4.

#### Incentivized Vigor Task

Participants performed a speeded precision joystick reaching task under three incentive conditions: jackpot ($1.60 reward), robber ($1.60 loss avoidance), or standard ($0.20 reward). The task comprised 300 trials (10% jackpot, 10% robber, 80% standard) across three runs. Success required reaching the cued target within a fixed deadline (1.87 s), calibrated from pilot data to yield ∼50% success. Behavioral analyses used repeated-measures ANOVA with factors of context, hold period, and run. Full task details are provided in SI Methods S5.

#### fMRI Preprocessing and Modeling

EPI data were preprocessed using ANTs and FSL, including motion correction, slice-time correction, fieldmap-based distortion correction, and normalization to MNI-152 space with 5 mm smoothing. First-level GLMs included instruction-phase regressors (stick functions at cue onset) and go-phase regressors (duration-scaled by RT) for each incentive context, plus nuisance regressors for hold period and reach execution. ROI-averaged parameter estimates were submitted to hierarchical Bayesian models with Student’s t likelihoods to accommodate outliers. Credible activation was determined by 89% highest-density intervals excluding zero. Full preprocessing, modeling, and inference details are provided in SI Methods S6.

## Acknowledgements

Authors thank Mario Mendoza at UCSB’s Brain Imaging Center and Katherin Martin, Arthur Toga, Danny Wang, Kay Jann, and Ioannis Pappas at the USC Mark and Mary Stevens Neuroimaging and Informatics Institute for scan acquisition development and data collection assistance. Authors also thank Logan Dowdle and Dan Handwerker (Tedana team) for acquisition development and Tedana preprocessing guidance. This research was funded in whole by Aligning Science Across Parkinson’s ASAP-020-519 through the Michael J. Fox Foundation for Parkinson’s Research (MJFF). For the purpose of open access, the authors have applied a CC BY public copyright license to all Author Accepted Manuscripts arising from this submission.

## Data Availability Statement

Anonymized data underlying this study will be deposited in a community-recognized, persistent repository and made publicly available no later than the time of publication of the final peer-reviewed article. The data, code, protocols, and key materials used and generated in this study are listed in a Key Resource Table alongside their persistent identifiers in the Supplementary Information (SI Table 4).

