## Supplementary material for "Incentive valence differentially engages open- and closed-loop basal ganglia circuits during movement initiation": SI

*Note: Clickable navigation links are provided in the footer of each page.*


**Table 1** Glossary of abbreviations

**Table 2** Regions of interest: atlas sources and definitions

**Table 3** Main behavior effects from repeated measures ANOVA

**Figure S1** Kinematic variables for each incentive context

**Figure S2** Connectome plot; seed in right PUTv

**Figure S3** Connectome plot; seed in right PUTd

**Figure S4** Connectome plot; seed in right PUTv (partial correlation\PUTd)

**Figure S5** Connectome plot; seed in left PUTd (partial correlation\PUTv)

**Figure S6** Connectome plot; seed in right PUTd (partial correlation\PUTv)

**Figure S7** Connectome plot; seed in left PUTv (partial correlation\BLA)

**Figure S8** Connectome plot; seed in right PUTv (partial correlation\BLA)

**Figure S9** Connectome plot; seed in left PUTv (partial correlation\CeA)

**Figure S10** Connectome plot; seed in right PUTv (partial correlation\CeA)

**SI Methods S1** Experiment 1: Participants and Study Protocol

**SI Methods S2** Experiment 1: Multi-Echo fMRI Preprocessing and Denoising with Tedana

**SI Methods S3** Experiment 1: CONN Functional Connectivity Analysis

**SI Methods S4** Experiment 2: Participants and Study Protocol

**SI Methods S5** Experiment 2: Incentivized Vigor Task

**SI Methods S6** Experiment 2: Task-Based fMRI Preprocessing, Modeling, and Bayesian Inference

**SI References** Shared bibliography for SI Methods

**Data & Code** Repositories for analysis code and datasets

**Table 4** Key Resource Table

Table 1 Glossary of abbreviations.

| **Abbreviation** | **Definition** |
|---|---|
| BLA | Basolateral amygdala |
| CeA | Central nucleus of the amygdala |
| CLC | Closed-loop circuit |
| CMA | Cingulate motor area |
| GPi | Globus pallidus internus |
| HDI | 89% highest density interval |
| fMRI | Functional magnetic resonance imaging |
| $M1_{UL}$ | Primary motor cortex, upper-limb region |
| $M1_{exUL}$ | Primary motor cortex, excluding upper-limb region |
| NAc | Nucleus accumbens |
| NBM | Nucleus basalis of Meynert |
| OLC | Open-loop circuit (putative) |
| PD | Parkinson's disease |
| PK | Paradoxical kinesia |
| PMd | Dorsal premotor cortex |
| PMv | Ventral premotor cortex |
| PUTd | Dorsal (sensorimotor) putamen |
| PUTv | Ventral (limbic) putamen |
| SA | Septal area of the basal forebrain |
| SMA | Supplementary motor area |
| SNpc | Substantia nigra pars compacta |
| VL | Ventrolateral thalamus |

Table 2 Regions of interest: atlas sources and definitions

| ROI | Atlas | Description | Reference |
|---|---|---|---|
| *Open-Loop Circuit / Ventral Striatum* | | | |
| BLA | Tyszka amygdala | BL masks combined (BL_BLV, BLN_BL_BLD+BLI, BLN_BM, BLN-La) | Tyszka & Pauli, 2016 |
| CeA | Tyszka amygdala | Central nucleus masks (CEN + CMN) | Tyszka & Pauli, 2016 |
| PUTv | Harvard-Oxford | Putamen ≥50%, voxels below MNI z = -1 | Di Martino et al., 2008 |
| NBM | JuBrain (SPM) | Sublenticular basal forebrain (Ch4) | Zaborszky et al., 2008 |
| SA | JuBrain (SPM) | Septum & horizontal diagonal band (Ch1-3) | Zaborszky et al., 2008 |
| NAc | Harvard-Oxford | Accumbens ≥50% | Desikan et al., 2006 |
| *Closed-Loop Circuit* | | | |
| PUTd | Harvard-Oxford | Putamen ≥50%, voxels above MNI z = -1 | Di Martino et al., 2008 |
| VL | Julich Brain | VLA + VLP, 207-area atlas at 50% | Amunts et al., 2020 |
| GPi | Tyszka subcortical | Deterministic atlas region | Pauli et al., 2018 |
| STN | Tyszka subcortical | Deterministic atlas region | Pauli et al., 2018 |
| *Motor & Premotor Cortex* | | | |
| $M1_{UL}$ | Brainnetome | Upper limb M1 ≥50% | Fan et al., 2016 |
| CMA | Hand-drawn | STG-defined using MNI coords, SMA reference | Paus, 2001 |
| SMA | Julich Brain | Area 6mp, 207-area atlas at 50% | Amunts et al., 2020 |
| PMd | Julich Brain | Areas 6d1-3, 207-area atlas at 50% | Amunts et al., 2020 |
| PMv | Julich Brain | Areas 6r1 + 6v1-3, 207-area atlas at 50% | Amunts et al., 2020 |

Note: All ROIs defined in MNI152 standard space. PUTv/PUTd threshold based on Talairach z = 2 converted to MNI z = -1.

Table 3 Main Effects from Repeated Measures ANOVA

| | | Effects | | | | | |
|---|---|---|---|---|---|---|---|
| | | Cue | | Run | | Hold | |
| Variable | Stat | F | p | F | p | F | p |
| success rate | mean | 4.2 | **.01** | 0.5 | .59 | 51.9 | **<.001** |
| RT | median | 18.5 | **<.001** | 0.3 | .75 | 110.5 | **<.001** |
| false starts | mean | 25.4 | **<.001** | 0.8 | .43 | 143.3 | **<.001** |
| max. velocity | median | 9.9 | **<.001** | 0.3 | .73 | 4.5 | **.03** |
| max. acceleration | median | 18.0 | **<.001** | 1.2 | .30 | 0.2 | .64 |
| time to max. vel. | median | 15.9 | **<.001** | 0.5 | .63 | 97.7 | **<.001** |
| time to max. acc. | median | 5.0 | **.007** | 0.2 | .79 | 61.3 | **<.001** |
| initial X position | median | 1.6 | .20 | 0.8 | .45 | 0.1 | .82 |
| initial Y position | median | 0.2 | .81 | 0.6 | .54 | 1.8 | .18 |
| time to quarter acc. | median | 18.6 | **<.001** | 0.7 | .49 | 111.6 | **<.001** |

Note: Bold values indicate $p < 0.05$. Degrees of freedom: Cue (2, error), Run (2, error), Hold (1, error).

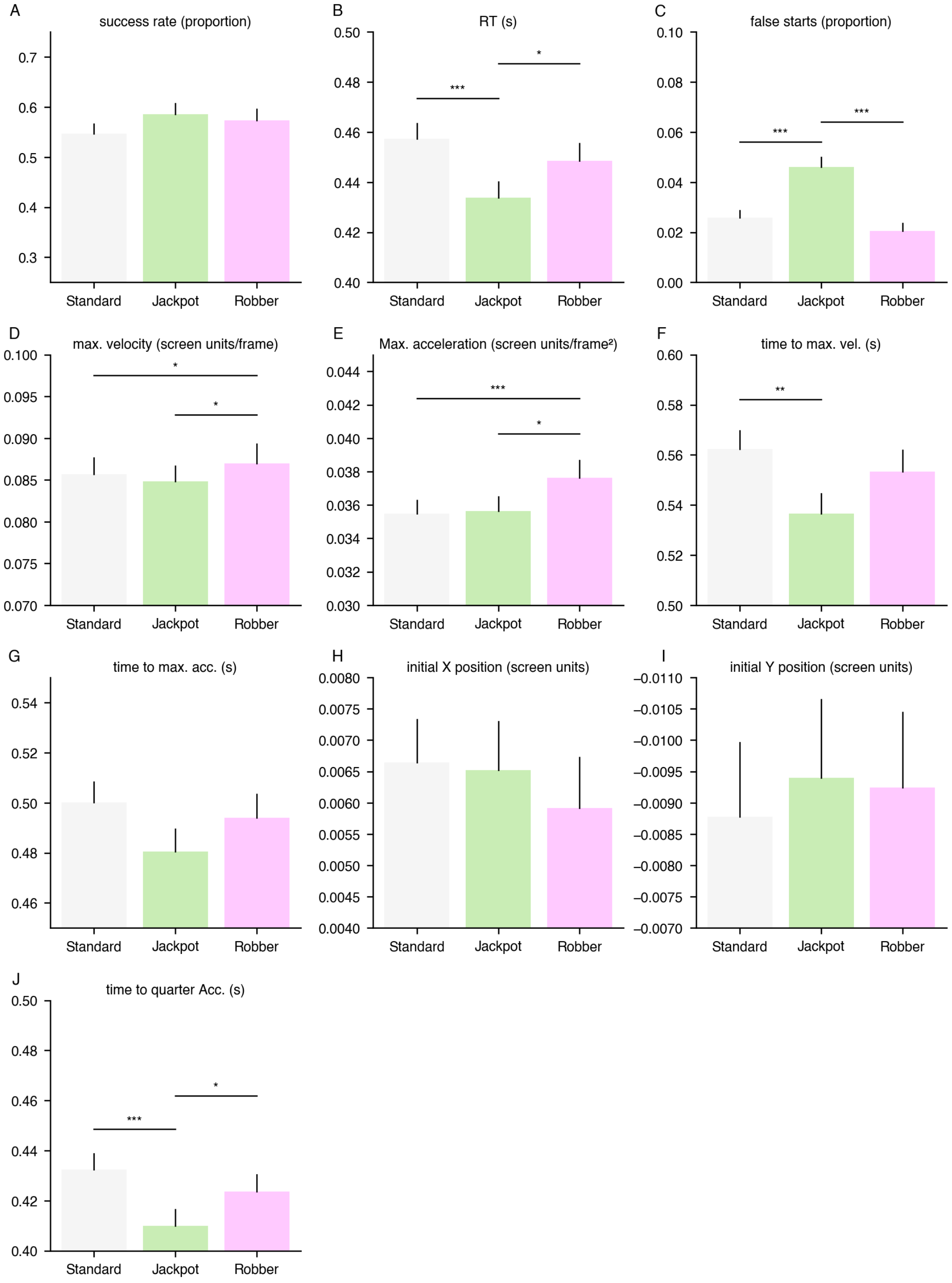


Figure S1 Kinematic variables for each incentive context. All measures are median per subject, and mean across subject, except proportion success and proportion false start. Figure annotated with pairwise comparison (FDR corrected) if main effect of cue significant from ANOVA; full results in SI Table 3

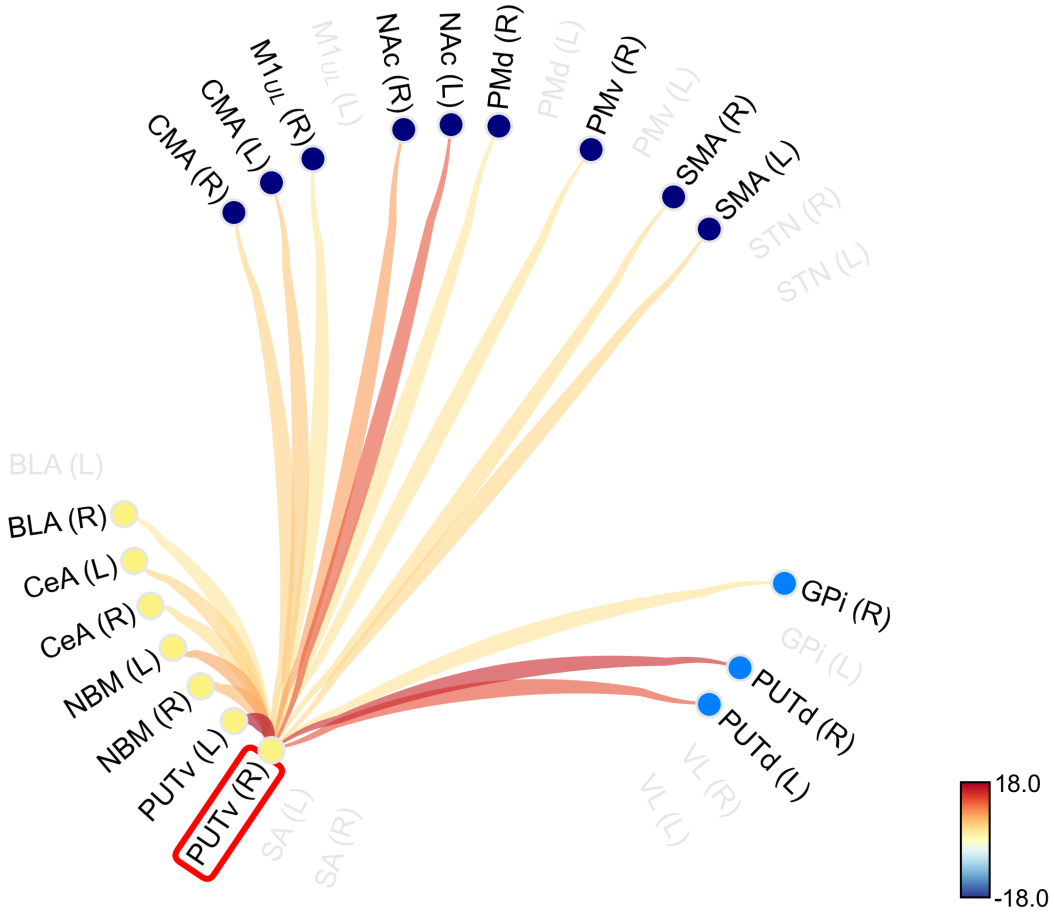


Figure S2 Connectome plot using seed in right ventral putamen

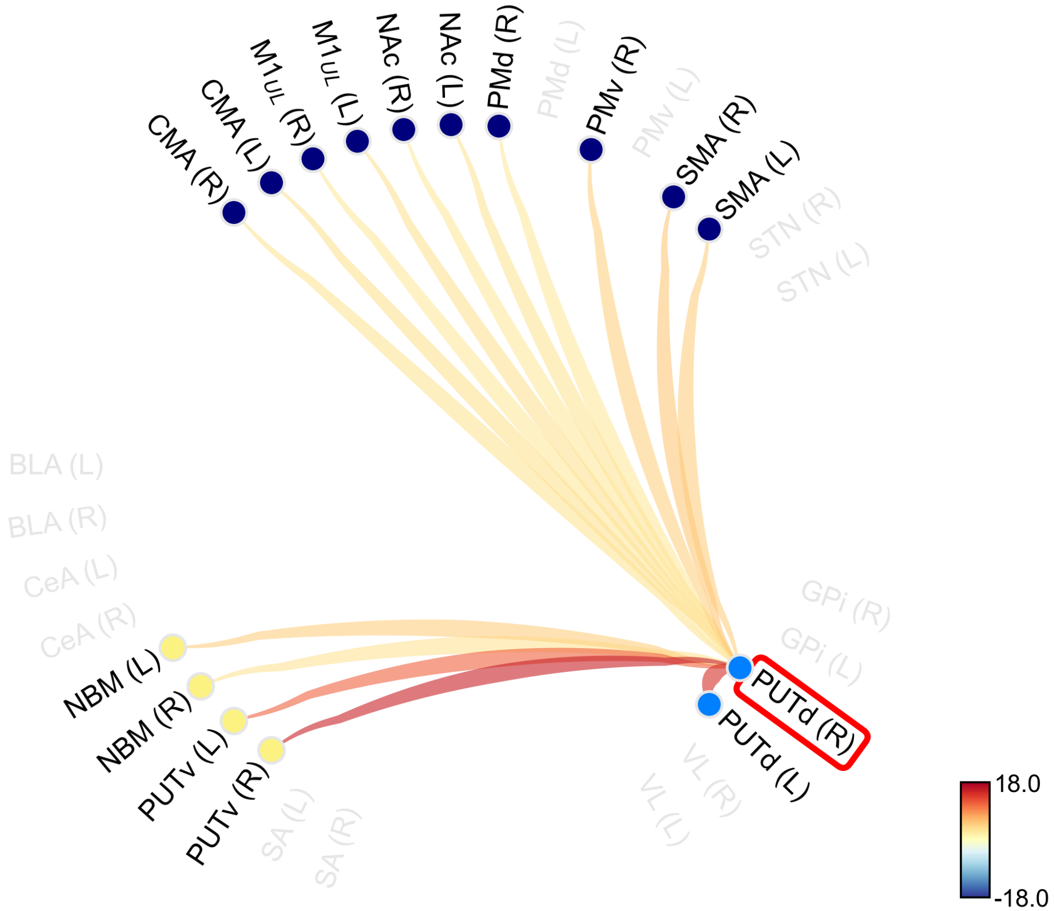


Figure S3 Connectome plot using seed in right dorsal putamen

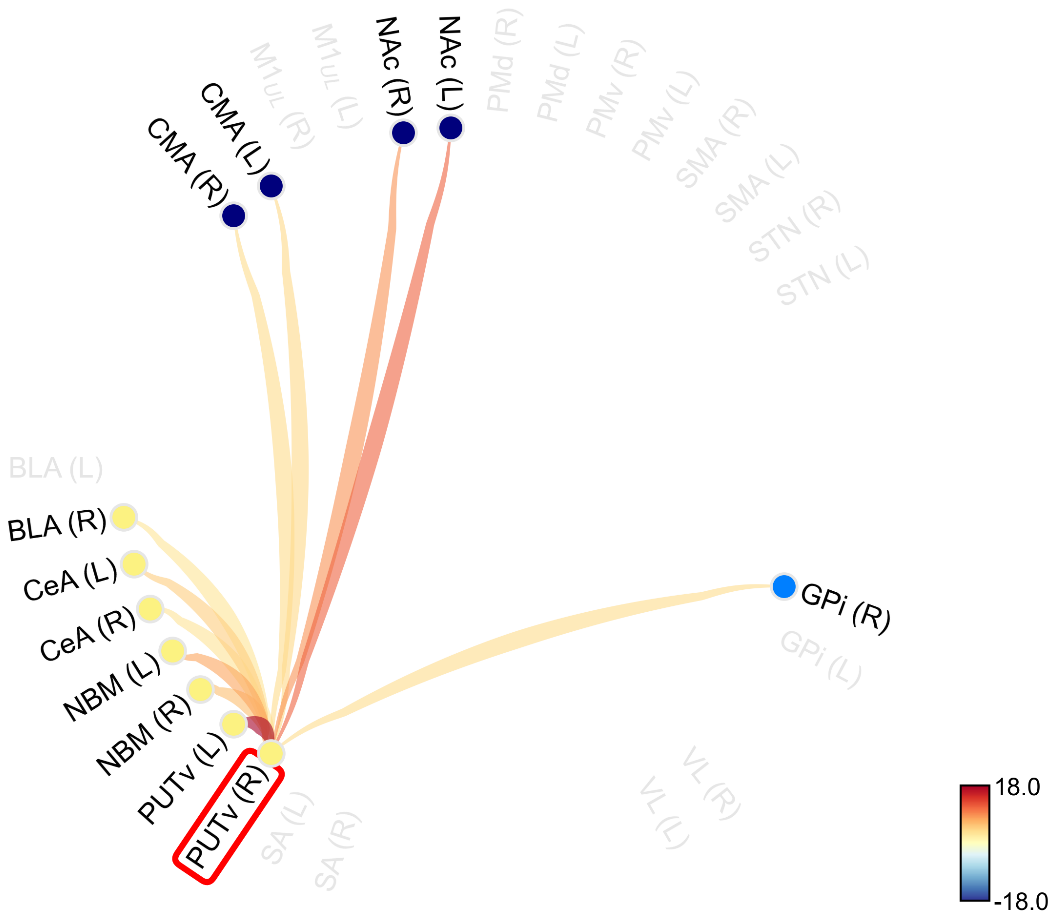


Figure S4 Connectome plot using seed in right ventral putamen; partial correlation (variance from dorsal putamen removed)

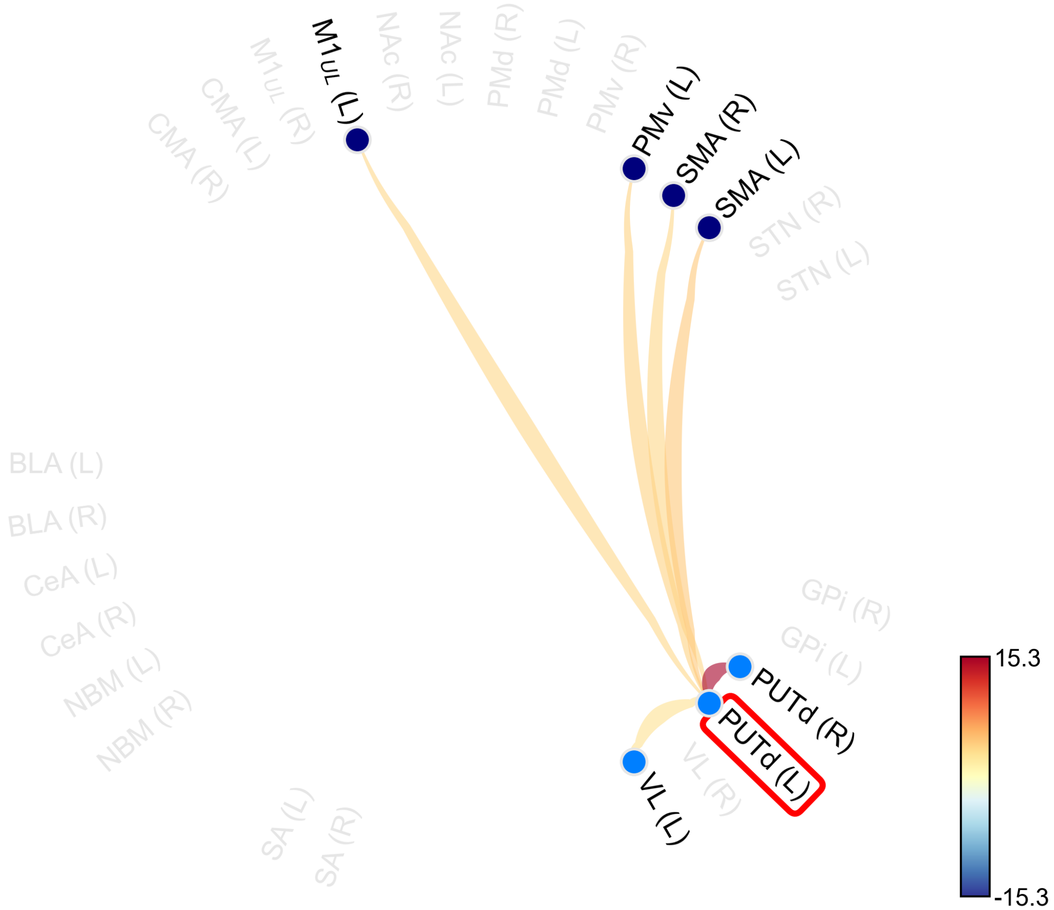


Figure S5 Connectome plot using seed in left dorsal putamen; partial correlation (variance from ventral putamen removed)

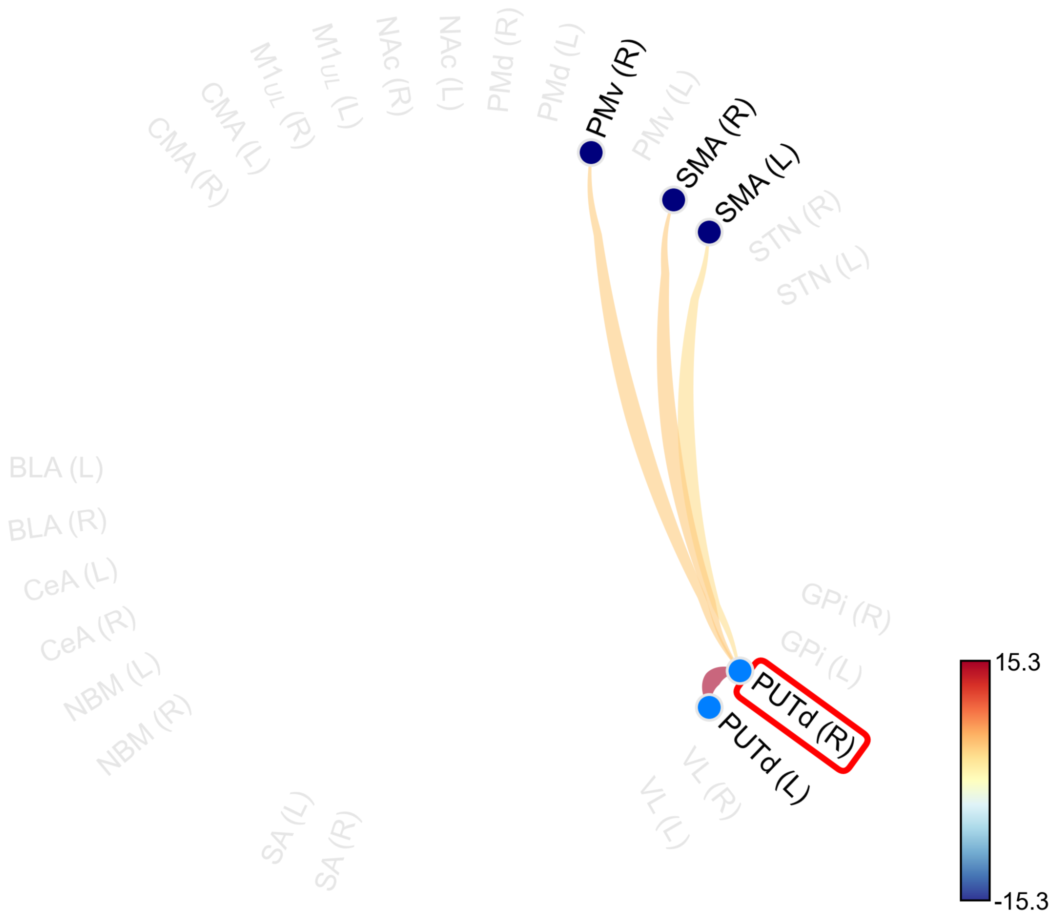


Figure S6 Connectome plot using seed in right dorsal putamen; partial correlation (variance from ventral putamen removed)

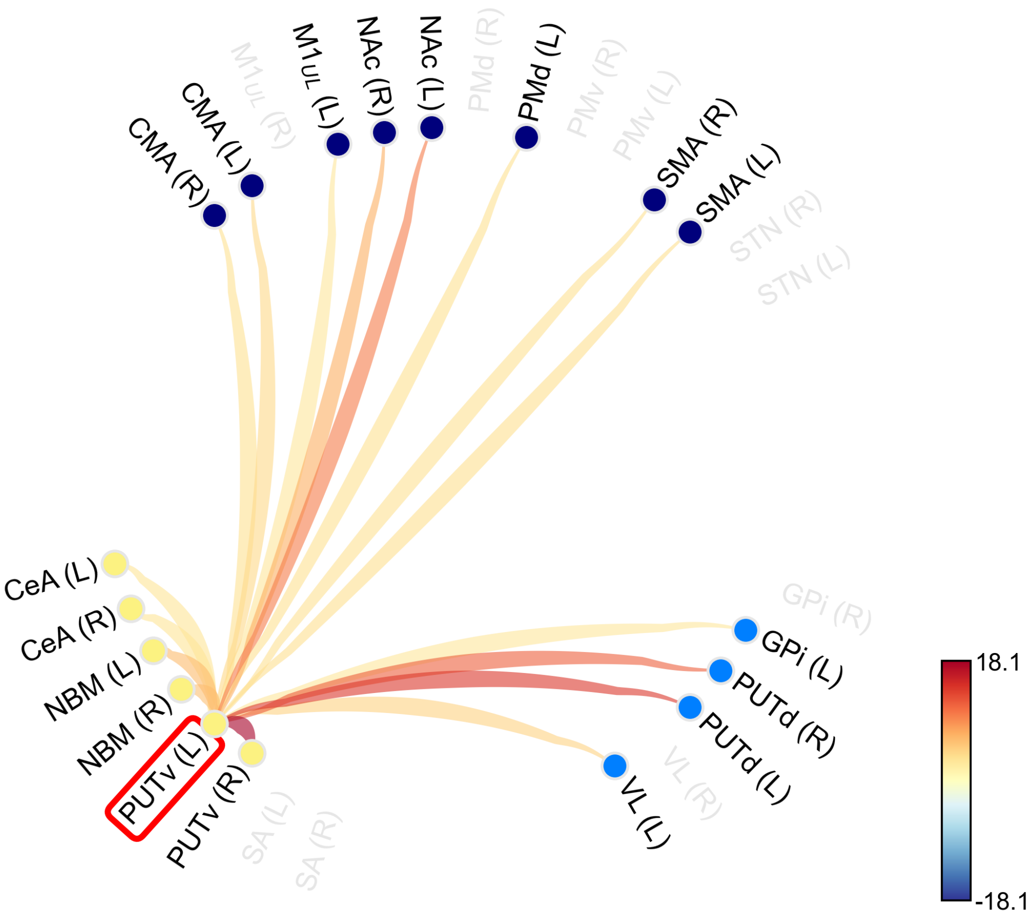


Figure S7 Connectome plot using seed in left ventral putamen; partial correlation (variance from basolateral amygdala removed)

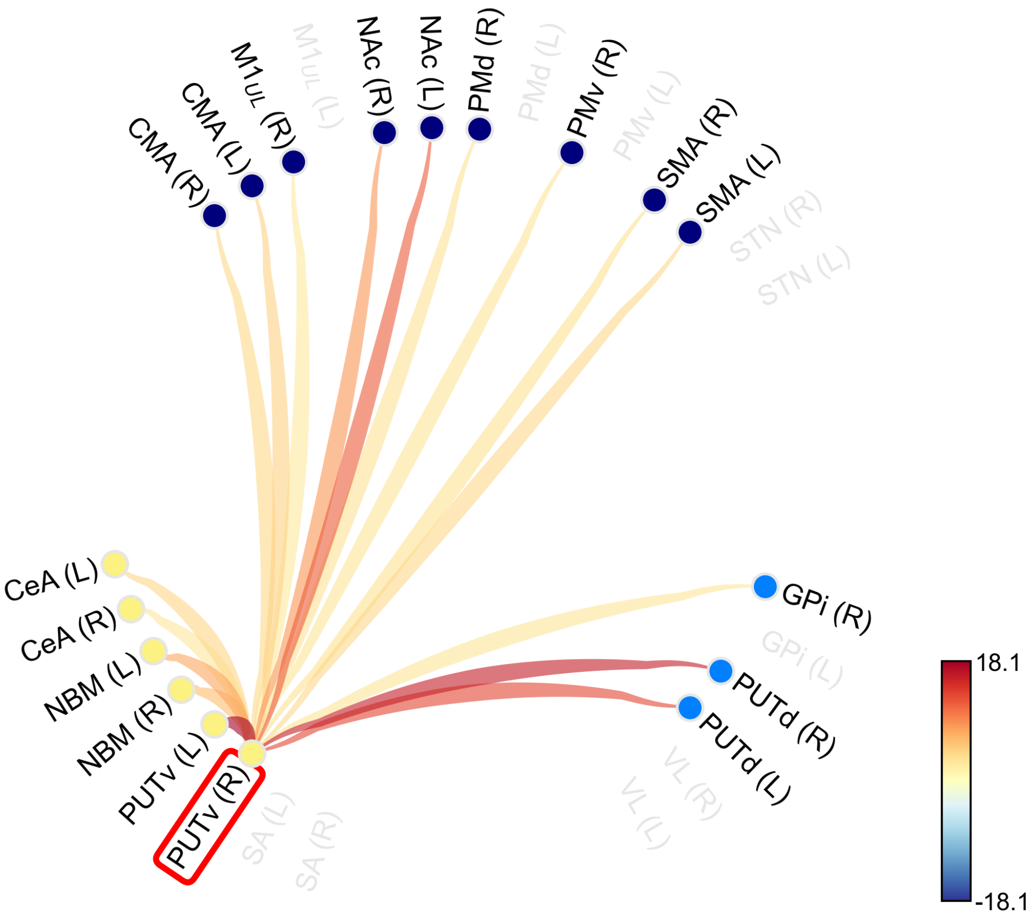


Figure S8 Connectome plot using seed in right ventral putamen; partial correlation (variance from basolateral amygdala removed)

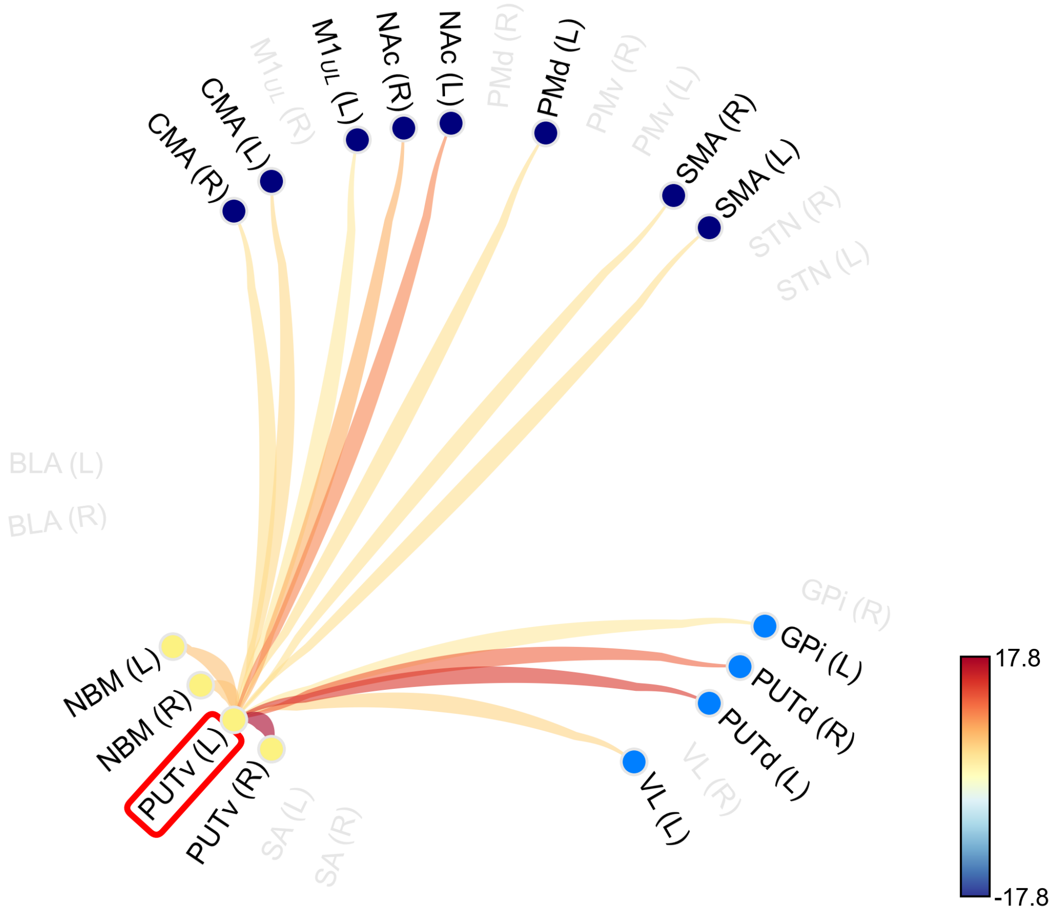


Figure S9 Connectome plot using seed in left ventral putamen; partial correlation (variance from central nucleus of amygdala removed)

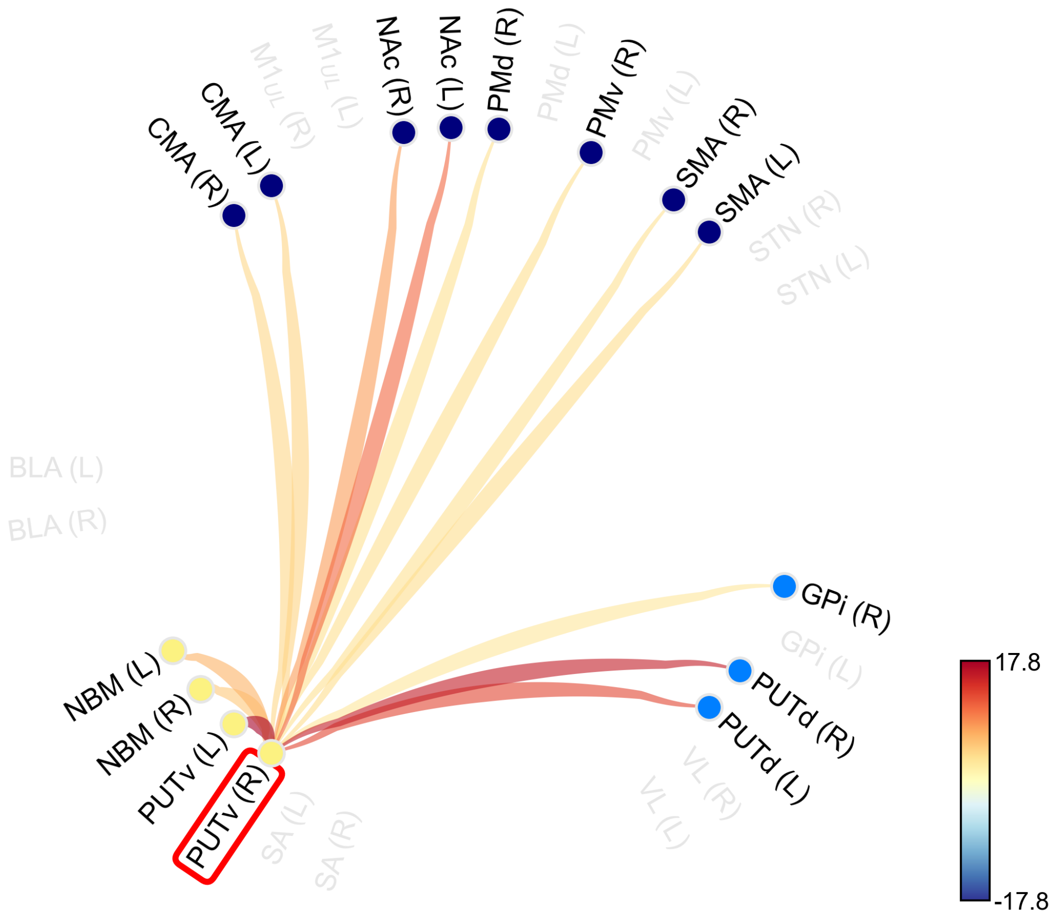


Figure S10 Connectome plot using seed in right ventral putamen; partial correlation (variance from central nucleus of amygdala removed)

### SI Methods

#### S1. Experiment 1: Participants and Study Protocol

A total of 28 young, healthy participants (15M/13F, mean age = 29.68 years; range = 21-39, all right-handed) completed all study recruitment, pre-screening, and protocol procedures. Study inclusion criteria required that participants be between the ages of 18 – 40 and competent with English enough to understand instructions. Exclusionary criteria were contraindications to MRI (non-removable metal, permanent mouth retainers/braces, incompatible medical devices, vertigo, dizziness, hearing loss/tinnitus, claustrophobia), pregnancy, or clinically significant cardiac, neurological, pulmonary, or psychiatric diseases. Participants were recruited via word of mouth and digital flyers sent to the University of California, Santa Barbara and University of Southern California communities. To assess eligibility, participants completed online screening forms that assessed demographics, health history, and MRI safety eligibility. All participants provided written informed consent for study procedures approved by the Institutional Review Boards at University of California, Santa Barbara and the University of Southern California and were paid 20 U.S. dollars/hour for the full MRI session. Data were acquired at the Center for Image Acquisition at the University of Southern California Mark and Mary Stevens Neuroimaging and Informatics Institute.

On the day of their study session, participants completed the Edinburgh Handedness Inventory (Oldfield, 2013) and a comprehensive questionnaire assessing demographics, medication use, and health history. Participants then completed a 30-minute MRI protocol at the University of Southern California's Center for Image Acquisition, where they were scanned with a Siemens 7T Terra scanner and a research-only 8-channel transmit 32-channel receive Nova 8Tx/32Rx head coil (Nova Medical Inc.) First, high-resolution T1-weighted magnetization prepared rapid gradient echo (MPRAGE) anatomical scans were acquired (TR = 4300 ms, TE = 2.27 ms, FOV = 240 mm, T1 = 1000 ms, flip angle = 4°, with 0.75 $mm^3$ voxel size). Following the anatomical scan, modified CMRR multi echo resting-state echo planar imaging (EPI) sequences were acquired (Total time = 10:52 mins, TR = 2100ms, flip angle = 70°, 1.5mm thick axial slices, 2x2 mm in plane, 300 volumes, 200mm FOV, iPat GRAPPA factor = 3, MB factor = 4, phase encoding = posterior to anterior), where participants were instructed to keep their eyes open. The echo times and slice number were slightly modified for the first 8 participants (TE1=15.20ms, TE2=34.23ms, TE3=53.26ms, 84 slices) vs. the remaining 13 (TE1=15.20ms, TE2=33.87ms, TE3=52.54ms, 100 slices). Two spin-echo EPI sequences for field map generation were also acquired: one with anterior to posterior (AP) and another with posterior to anterior (PA) phase encoding to use for distortion correction (TR = 3000ms, TE = 60ms, flip angle = 70°, 1.5mm thick axial slices, 2x2mm in plane, 192mm FOV, iPat GRAPPA factor = 2, MB factor = 5). During scanning, participant respiration and pulse measurements were acquired with an in-scanner Siemens respiration belt and pulse sensor for later denoising.

#### S2. Experiment 1: Multi-Echo fMRI Preprocessing and Denoising with Tedana

We conducted MRI preprocessing with a custom pipeline featuring Advanced Normalization Tools (ANTs) (Avants et al., 2011), FSL (Jenkinson et al., 2012), and TE Dependent ANAlysis version 24.0.1 (DuPre et al., 2021). Our preprocessing pipeline was inspired by Lynch et al., 2020. First, T1-weighted anatomical data were skull-stripped with antsBrainExtraction.sh and resampled to functional image resolution (2mm voxels). We then used FSL Topup with AP and PA spin echo sequences to obtain an undistorted magnitude image and fieldmap for distortion correction. Next, we averaged the EPI data single band reference images at each timepoint across 3 echoes; this reference image was then brain extracted with FSL BET. EPI data were motion-corrected with FSL MCFLIRT (transforms for echo 1 were applied to echoes 2 and 3) and slice time corrected with FSL slicetimer. We used FSL's epi reg with the single-band reference brain to obtain fieldmap distortion correction transforms and co-registration transforms to T1-MPRAGE data. These transforms were then applied to EPI data to place all three echoes into participant-specific anatomical space. Due to fieldmap scan acquisition error, the distortion correction transforms were not applied to the first 12/28 participants.

Next, we denoised resting-state EPI data in anatomical space with Tedana (see Tedana program output below for full details). Briefly, Tedana optimally combines the three echo images, reduces data dimensionality with principal component analysis (PCA), and removes TE-independent (non-BOLD) components from the data with independent component analysis (ICA). After Tedana denoising, we spatially normalized all participant data by registering participant T1-anatomical images to the MNI-152 2mm template and applying this transformation to the denoised EPI data. Normalized EPI data were then inputted into the CONN Toolbox (Whitfield-Gabrieli et al., 2012; see S3. CONN Functional Connectivity Analysis).

**Tedana program output:**

Following standard preprocessing, multi-echo EPI data were denoised using TE-dependent analysis (tedana workflow v24.0.1). The ICA decision tree employed was tedana_orig, structurally analogous to the MEICA v2.5 criteria described by Kundu et al. (2013) and detailed by Olafsson et al. (2015). A user-defined mask was initially applied, after which an adaptive mask was generated using the dropout method. In this approach, each voxel's value reflects the number of echoes containing valid signal. A two-stage masking procedure was then implemented: a liberal mask (voxels with signal in at least the first echo) was used for optimal combination, T2*/S0 estimation, and denoising, while a conservative mask (restricted to voxels with signal in at least the first three echoes) was applied during component classification.

A monoexponential decay model was fit to each voxel's signal using nonlinear optimization to estimate T2* and S0 maps, with initial values derived from log-linear fits. The adaptive mask determined which echoes contributed to each voxel's parameter estimation. When nonlinear fitting failed, log-linear estimates were retained. Multi-echo data were optimally combined using the T2*-weighted method (Posse et al. 1999). Global signal regression was applied to both multi-echo and optimally combined datasets before dimensionality reduction.

Principal component analysis with pre-specified component count was applied to the optimally combined data. The following metrics were computed at this stage: kappa (TE-dependence), rho (TE-independence), countnoise, countsigFT2, countsigFS0, dice_FT2, dice_FS0, signal-noise_t, variance explained, normalized variance explained, and d_table_score. Kappa and rho quantify the degree to which components track T2* decay (BOLD-like) versus remain invariant across TEs (non-BOLD artifacts). A t-statistic (signal-noise_z) and associated p-value (signal-noise_p) were derived by contrasting T2*-model F-statistics between cluster voxels (signal) and non-cluster voxels (noise), measuring component association with signal over noise. The count of significant non-cluster voxels was also tallied per component.

Independent component analysis then decomposed the dimensionally reduced data. Component-level metrics identical to those from PCA were recomputed on ICA outputs. Component classification proceeded via the tedana_orig decision tree to identify BOLD (TE-dependent) versus non-BOLD (TE-independent) sources. Rejected components were removed from the optimally combined data, yielding denoised time series for subsequent connectivity analyses. All computations utilized numpy (Van Der Walt et al. 2011), scipy (Virtanen et al. 2020), pandas (McKinney 2010; pandas development team 2020), scikit-learn (Pedregosa et al. 2011), nilearn, nibabel (Brett et al. 2019), matplotlib (Hunter 2007), and bokeh (Bokeh Development Team 2018). Dice similarity indices were computed as described by Dice (1945) and Sorensen (1948).

#### S3. Experiment 1: CONN Functional Connectivity Analysis

Following tedana denoising and PhysIO-based physiological noise removal, residual confounds were addressed using the CONN functional connectivity toolbox (release 22.a; Nieto-Castanon 2022). Anatomical noise components were extracted via CompCor (Behzadi et al. 2007) by computing the five largest principal components orthogonal to the mean BOLD signal within each subject's eroded white matter mask, plus the first principal component from eroded CSF. These were combined with the 18 RETROICOR regressors already applied and additional linear trends within each run. Temporal filtering applied a bandpass of 0.008–0.09 Hz to the denoised BOLD time series, following recommendations by Hallquist et al. (2013) to avoid reintroducing noise via spectral misspecification. The effective degrees of freedom post-denoising, accounting for all confound regressors, were estimated to range from 60.6 to 92.3 (mean 88.0) across participants (Nieto-Castanon, submitted).

Seed-to-voxel connectivity was characterized using bivariate correlation coefficients estimated via weighted general linear model (Nieto-Castanon 2020a). To compensate for transient magnetization at run onset, individual volumes were weighted by a step function convolved with the SPM canonical hemodynamic response and rectified. Fisher-transformed correlation maps were generated for each seed ROI. ROI-level inference combined connection-level statistics across all voxels projecting from each seed using multivariate parametric tests with random effects across subjects (Nieto-Castanon 2020b). Significant connections were identified using false discovery rate correction (Benjamini-Hochberg procedure) applied across the upper triangle of the connectivity matrix to avoid double-counting symmetric connections, following the recommended CONN toolbox approach. The FDR threshold was set at $\alpha = 0.05$. Connection strengths are reported as t-statistics from the correlation analysis (Benjamini & Hochberg 1995; Nieto-Castanon 2020c).

Partial correlation analyses isolated PUTv-specific connectivity by regressing the ipsilateral PUTd mean time series from the PUTv seed prior to connectivity estimation. This orthogonalization removed shared variance attributable to canonical motor circuitry. Seed-to-voxel maps were recomputed using the residualized PUTv signal, with identical statistical thresholding applied. Network-level comparisons extracted mean connectivity values across all voxels within anatomically defined target ROIs. Motor network ROIs comprised CMA, SMA, PMd, $\mathrm{M1_{UL}}$, and PMv; limbic network ROIs comprised BLA, CeA, and NBM. Repeated-measures ANOVA tested for putamen subregion by target network interactions, with Bonferroni correction applied to post-hoc pairwise comparisons. All analyses were conducted in CONN (RRID:SCR_009550) and SPM12 (v7487; Wellcome Centre for Human Neuroimaging).

#### S4. Experiment 2: Participants and Study Protocol

A total of 68 young, healthy participants (50F/18M, mean age = 20.75 years; SD = 1.86) completed all study recruitment, pre-screening, and protocol procedures. Study inclusion criteria required that participants be between the ages of 18–40 and competent with English enough to understand instructions. Exclusionary criteria were contraindications to MRI (non-removable metal, permanent mouth retainers/braces, incompatible medical devices, vertigo, dizziness, hearing loss/tinnitus, claustrophobia), pregnancy, or clinically significant cardiac, neurological, pulmonary, or psychiatric diseases. Participants were recruited via word of mouth and a study recruitment portal at the University of California, Santa Barbara. To assess eligibility, participants completed online screening forms that assessed demographics, health history, and MRI safety eligibility. All participants provided written informed consent for study procedures approved by the Institutional Review Board at University of California, Santa Barbara.

Each subject's data were collected in a single session, lasting approximately two hours. fMRI data were recorded using a Siemens 3T Prisma scanner with a 64-channel phased-array head/neck coil. Task responses were recorded using an MR-compatible joystick. This was fixed to the center of a wooden table, which was placed across participants' torsos once they were supine in the scanner bore. The position of the table was adjusted along the bore to ensure a comfortable distance for the participant's arm. Arms were supported by pillows and padding.

Each session began with a T1-weighted magnetization prepared rapid gradient echo (MPRAGE) anatomical scan (TR = 2500ms, TE = 2.22ms, FOV = 241mm, T1 = 851ms, flip angle = 7°, $0.94mm^3$ voxel size). We next acquired a double-echo gradient echo field map sequence for distortion correction (TR = 758 ms, TE = 4.92ms, flip angle = 60°, $2.5mm^2$ thick axial slices). We then acquired 4-dimensional echo-planar imaging during each of the three runs of the task (TR = 1900 ms, TE = 30 ms, flip angle = 65°, $2.5mm^3$ voxel size, multiband acceleration factor of 2).

#### S5. Experiment 2: Incentivized Vigor Task

Participants performed a speeded precision joystick reaching task in which they moved a cursor to a cued target under one of three incentive conditions: jackpot ($1.60 reward), robber ($1.60 loss avoidance), or standard ($0.20 reward). The task comprised 300 trials (10% jackpot, 10% robber, 80% standard) across three runs.

The deadline for reach completion from onset of the go cue (1.87s) was calibrated from a pilot sample to achieve a 50% success rate across participants. This pilot sample of eight participants performed the task inside the scanner environment. From their data we used Bayesian estimation of the group-level mean and standard deviation of log-transformed completion times to infer the time associated with a 50% probability of completion.

Visual stimuli were rear-projected onto a screen placed approximately 110cm behind participants using an LCD projector (1920×1080, 60 Hz). Participants viewed rectified images by way of a double mirror in the head coil. Each trial began with participants holding a screen cursor (radius=0.5°; yoked to joystick position) at a starting position (radius=1.2°; ∼9.5° below screen center). An instruction cue then appeared at one of two spatial locations. Each location had a radius of 1.2° and were both ∼9.5° above the screen center. The left target was $\sim 19^\circ$ to the left of the screen's midline, and the right target $\sim 19^\circ$ to the right. Each instruction cue was made of the same small blue squares, re-arranged to make a neutral (standard trial), happy (jackpot) or sad (robber) face. Each stimulus was $\sim 6.6^\circ$ radius and appeared centered on one of the two spatial targets for 200 ms. The hold period prior to the instruction cue (either 1 or 2s) and go cue (1 - 4s) altered across trials according to an m sequence. Total completion time on each trial was the time from the onset of the go cue until they reached the cued target, holding it in place for an additional 0.8s. Completions within the deadline (1.87s) were successful, with verbal feedback indicating the trial's reward alongside the word "success". Completions slower than this deadline were unsuccessful, with verbal feedback indicating the trial's loss alongside the word "too slow". All text was $\sim 2^\circ$ in height and appeared at the screen center. Participants performed a set of 40 training trials prior to the experimental runs, where difficulty gradually increased from a deadline of 5s to the task target.

Cursor position was continuously recorded on each frame. Movement initiation time (RT) was defined as the time taken to initiate movement (cursor moving outside starting position, i.e., centroid exceeding radius of the starting position) relative to the onset of the go-cue. False-starts were any movement of the cursor outside of the starting position after the onset of the instruction cue, prior to the onset of the go cue. Behavioral analyses of success rate, reaction time, false starts, maximum velocity, maximum acceleration, time to maximum velocity, time to maximum acceleration, initial cursor position, and time to quarter acceleration were conducted with repeated-measures ANOVA using factors of context (standard, jackpot, robber), hold period ($>3$ s,$\leq 3$s) and run (1,2,3). Post-hoc pairwise correction for multiple comparison used FDR.

#### S6. Experiment 2: Task-Based fMRI Preprocessing, Modeling, and Bayesian Inference

We conducted MRI preprocessing with a custom pipeline featuring Advanced Normalization Tools (Avants et al., 2011) and FSL (Jenkinson et al., 2012). T1-weighted anatomical data were skull-stripped with antsBrainExtraction.sh and resampled to functional image resolution (2mm voxels). We then applied motion, slicetime, and distortion correction with field maps to EPI data with FSL FEAT. Next, we used ANTs (antsRegistrationSyN) to coregister and apply a non linear transform of EPI data to respective T1 anatomical data, then to MNI152 2mm space. EPI data were smoothed with a 5 mm FWHM Gaussian kernel.

First-level GLMs were fitted to preprocessed EPI data using FSL FEAT. The model included seven regressors of interest plus two nuisance regressors. Instruction-phase regressors were modeled as stick functions (unit height, 0.1 s width) event-locked to cue onset, separately for standard, jackpot, and robber trials. Go-phase regressors were duration-scaled and event-locked to go cue onset, also separated by context. For a trial with initiation time (RT) of $t$ seconds, the corresponding go-phase regressor had duration $t$ and unit height. Under this parametrization, slower initiation (longer RT) produces longer regressor duration, such that positive regression coefficients indicate greater BOLD signal with slower movement initiation. For interpretability, plotted values are sign-reversed so that positive values reflect greater BOLD activity associated with faster initiation.

Nuisance regressors captured task-related activity not specific to RT effects. The pre-go hold regressor had onset at instruction cue offset and duration equal to the hold period (1–4 s, varied by trial). The reach execution regressor had onset at movement initiation (RT from go cue) and duration from initiation to trial end. Both nuisance regressors were unit height and duration-scaled. Additional confound regressors for motion parameters and temporal derivatives were included following standard FSL procedures.

Run-level parameter estimates (cope files) for each regressor were extracted and averaged within subjects to create subject-level summaries. For each ROI, mean activation across all voxels was computed per subject and regressor, yielding a single value per subject, per ROI, per contrast. These ROI-averaged values were then submitted to group-level Bayesian inference.

Hierarchical Bayesian models were implemented to estimate group-level activation parameters while accounting for between-subject variability and accommodating outliers. For each ROI and contrast combination, the distribution of subject-level activations was modeled as a Student's $t$ distribution with location parameter $\mu$, scale parameter $\sigma$, and degrees-of-freedom parameter $\nu$. A single mixture model was fitted across all ROI-contrast pairs simultaneously. Uninformed priors were assigned: $\mu_{\text{ROI,contrast}} \sim \mathcal{N}(0,1)$, $\sigma_{\text{ROI,contrast}} \sim \text{HalfNormal}(1)$, and a single $\nu \sim \text{HalfNormal}(1)$ was shared across all distributions to pool information about outlier prevalence.

Posterior distributions were sampled using Markov Chain Monte Carlo (MCMC) via the PyMC library. Four chains were run for 2000 tuning iterations followed by 2000 sampling iterations. Convergence was assessed via $\hat{R} < 1.01$ for all parameters. Circuit-level estimates were derived by averaging node-level posterior draws at each MCMC iteration: for circuit $c$ in context $x$, $\mu_{x,c} = \frac{1}{N}\sum_{i=1}^{N}\mu_{x,i}$ where $i$ indexes the $N$ nodes in circuit $c$. This approach propagates uncertainty from node-level to circuit-level estimates. Activations were deemed credibly nonzero when the 89% highest-density interval (HDI) of the posterior distribution excluded zero. Contrasts between circuits or contexts were evaluated by computing posterior differences ($\Delta = \mu_1 - \mu_2$) at each MCMC draw and assessing whether the resulting HDI excluded zero.

All Bayesian analyses were conducted in Python 3.11 using PyMC v5.10, ArviZ v0.17 for diagnostics, and NumPy v1.26 for numerical operations. Full model specifications and convergence diagnostics are available in the analysis code repository.

#### Data, Materials, and Software Availability

All raw imaging data are available open access in BIDS format through OpenNeuro as part of the SoCal Kinesia project (`https://www.socalkinesia.org`). Task paradigm code is available at `https://github.com/dundonnm/incentivized-vigor-task` and analysis scripts are available at `https://github.com/ejrise/incentive-valence`. All study resources are listed in Table 4 on the following page.

#### Table 4 Key Resource Table

| RESOURCE TYPE | RESOURCE NAME | SOURCE | IDENTIFIER | NEW or REUSE | ADDITIONAL INFORMATION |
|---|---|---|---|---|---|
| Dataset | SKIP highresFC MRI dataset | OpenNeuro | https://doi.org/10.18112/openneuro.ds005264.v1.1.0 | new | Raw Experiment 1 7T Resting MRI data |
| Dataset | SKIP incentivized reaching MRI dataset | OpenNeuro | https://doi.org/10.18112/openneuro.ds005263.v1.0.0 | new | Raw Experiment 2 3T Task MRI data |
| Protocol | 7T MRI protocol | protocols.io | dx.doi.org/10.17504/protocols.io.dm6gpbbj1lzp/v1 | new | Experiment 1 scanning protocol |
| Software/code | Incentivized Vigor Task | Github/Zenodo | https://doi.org/10.5281/zenodo.8216235 | new | Experiment 2 MRI Task code |
| Software/code | Analysis code and mask repo | Github | https://github.com/ejrise/incentive-valence | new | All Exp 1+2 brain masks and code |
| Software/code | FSL 6.0.7.12 | http://www.fmrib.ox.ac.uk/fsl/ | RRID:SCR_002823 | reuse | |
| Software/code | CONN Toolbox | https://web.conn-toolbox.org/ | RRID:SCR 009550 | reuse | |
| Software/code | TE Dependent ANAlysis (Tedana) 24.0.1 | https://tedana.readthedocs.io/en/stable/# | N/A | reuse | |
| Software/code | Advanced Normalization Tools (ANTs) 2.3.4 | https://github.com/ANTsX/ANTs | RRID:SCR_004757 | reuse | |
| Software/code | PhysIO toolbox 6.0.1 | https://www.nitrc.org/projects/physio/ | RRID:SCR_003430 | reuse | |
| Software/code | Python 3.11.5 | https://www.python.org/downloads/release/python-360/ | RRID:SCR_008394 | reuse | |
| Software/code | PyMC v. 5.10 | https://www.pymc.io/welcome.html | RRID:SCR_018547 | reuse | |
| Software/code | MATLAB | https://www.mathworks.com/products/matlab.html | RRID:SCR_001622 | reuse | |
| Other | 7T MRI Scanner | Siemens | MAGNETOM Terra | reuse | |
| Other | 3T MRI Scanner | Siemens | MAGNETOM Prisma | reuse | |
